# Transcription factors instruct DNA methylation patterns in plant reproductive tissues

**DOI:** 10.1101/2025.02.21.639562

**Authors:** Guanghui Xu, Yuhan Chen, Fuxi Wang, En Li, Julie A. Law

**Affiliations:** Plant Molecular and Cellular Biology Laboratory, Salk Institute for Biological Studies, La Jolla, CA, 92037, USA; Division of Biological Sciences, University of California, San Diego, La Jolla, CA 92093

**Keywords:** DNA methylation, siRNAs, REM transcription factors, RNA-directed DNA methylation, CLSY3

## Abstract

DNA methylation is maintained by forming self-reinforcing connections with other repressive chromatin modifications, resulting in stably silenced genes and transposons. However, these mechanisms fail to explain how new methylation patterns are generated. In Arabidopsis, CLASSY3 (CLSY3) targets the RNA-directed DNA methylation (RdDM) machinery to different loci in reproductive tissues, generating distinct epigenomes via unknown mechanism(s). Here, we discovered that several different REPRODUCTIVE MERISTEM (REM) transcription factors are required for methylation at CLSY3 targets specific to male or female reproductive tissues. We designate these factors as REM INSTRUCT METHYLATION (RIMs) and demonstrate that disruption of their DNA binding domains, or the motifs they recognize, blocks RdDM. These findings not only reveal RIMs as the first sex-specific RdDM proteins but also establish a critical role for genetic information in targeting DNA methylation. This novel mode of targeting expands our understanding of how methylation is regulated to include inputs from both genetic and epigenetic information.

## Introduction

DNA methylation is a conserved epigenetic modification that plays important roles in regulating gene expression and silencing transposons in both plants and animals^1^. A critical feature of epigenetic regulation is its capacity to facilitate two essential but opposing functions: the ability to maintain heritable patterns of chromatin modifications during cell division, while retaining the flexibility to generate diverse patterns during development or in response to the environment^2,3^. Over the last several decades research elucidating self-reinforcing connections between DNA methylation and histone modifications has provided key insights into the inheritance of DNA methylation, revealing how stable patterns of methylation are maintained^4^. However, it remains unclear how DNA methylation pathways are modulated to generate the distinct epigenetic profiles observed during plant development, regeneration, and reproduction^5–7^.

In the plant model *Arabidopsis thaliana*, DNA methylation patterns are dictated by interconnected pathways controlling the establishment, maintenance, and removal of methylation at cytosine bases^4,8–10^. DNA methylation in all three sequence contexts (CG, CHG and CHH, where H=A, T, or C) is initially established via the RNA-directed DNA methylation (RdDM) pathway^8^. This pathway employs two plant-specific RNA polymerases (Pol-IV and Pol-V) to generate 24nt small interfering RNAs (siRNAs) and long-noncoding RNAs, respectively^11–15^. Together, these RNAs facilitate the targeting of the de novo methyltransferase, DOMAINS REARRANGED METHYLTRANSFERASE 2 (DRM2), to transposons and repeats throughout the genome^16,17^. However, efficient targeting of both Pol-IV and Pol-V rely on chromatin modifications that are maintained by other DNA methylation pathways^4^. For example, Pol-IV targeting is controlled by a family of four SNF2-like remodeling factors, CLASSY (CLSY1-4)^18,19^, and loci regulated by CLSY1 and CLSY2 rely on SAWADEE DOMAIN HOMOLOG 1 (SHH1)^18^, a histone reader that recognizes H3K9 methylation^20,21^. However, H3K9 methylation is not deposited as part of the RdDM pathway. Instead, it is established and maintained by two other DNA methylation pathways that are interconnected with H3K9 methylation via self-reinforcing loops. These pathways are comprised of two families of dual reader-writer proteins^4,22^. The first family, SU(VAR)3-9 HOMOLOG 4 (SUVH4), SUVH5 and SUVH6, reads (*i.e.* binds to) DNA methylation^23,24^ and writes (*i.e.,* deposits) H3K9 methylation^25–27^. The second family, CHROMOMETHYLTRANSFERASE 2 (CMT2) and CMT3, have the opposite specificity, they read H3K9 methylation^28,29^ and write CHH and CHG DNA methylation^29–32^, respectively. Like Pol-IV, Pol-V targeting also depends on epigenetic modifications controlled by other DNA methylation pathways. Specifically, its targeting relies on the methyl-DNA binding activities of SUVH2 and SUVH9^33^, which read CHH methylation deposited by the RdDM pathway, CHG methylation maintained by the CMT3-SUVH pathway^34^, and CG methylation maintained by METHYLTRANSFERASE 1 (MET1)^34^. Together, these and other connections between DNA methylation pathways^4^ ensure that once established DNA methylation patterns are efficiently maintained, resulting in a stably repressed epigenetic state.

Beyond the local interactions connecting and reinforcing chromatin modifications, the activities of the different DNA methylation pathways are also regulated by chromatin on a global scale. While RdDM acts mainly at short transposons and repeats located in the chromosome arms, CMT2 and CMT3 act mainly at longer transposons and repeats enriched in pericentromeric heterochromatin^29,30^. This compartmentalization of pathways is associated with differences in the levels of two histone variants that influence chromatin compaction, H1 and H2A.W^4^. Low levels of H1 and H2A.W in the chromosome arms favor activity by RdDM while high levels, especially of H1, in pericentromeric heterochromatin suppresses RdDM^35–37^. For the CMT pathways, the suppressive effect of H1 is overcome by the activity of a chromatin remodeler, DECREASED DNA METHYLATION 1 (DDM1), resulting in higher activity in pericentromeric heterochromatin^30^. The boundaries between heterochromatin and euchromatin are also reinforced by several related DNA demethylases that act at these regions to prevent hypermethylation^38^.

Taken together, the field’s understanding of these DNA methylation pathways supports the long-standing dogma that DNA methylation patterns are regulated by chromatin features (*i.e.* self-reinforcing loops involving chromatin modifications, histone variants, and chromatin compaction levels) rather than by specific DNA features (*i.e.* sequence motifs). Nevertheless, current models fail to explain the tissue-specific targeting of the RdDM machinery to loci regulated by CLSY3, including the recently identified (and essentially non-overlapping) HyperTE^39^ and siren loci^40,41^ that dominate the siRNA landscapes in male (anther) and female (ovule) reproductive tissues, respectively. Unlike loci regulated by CLSY1 and CLSY2, siRNA production at loci controlled by CLSY3 and CLSY4 are not dependent on H3K9 methylation or non-CG methylation^18^, demonstrating they are not subject to the same self-reinforcing loops as other RdDM targets. Furthermore, as HyperTE and siren loci show very low methylation levels across other profiled tissues, it remains unclear how CLSY3 is targeted to such distinct sets of naïve loci in anthers and ovules tissues. One possible new mechanism for CLSY3 targeting was revealed by the identification of a conserved sequence motif at siren loci bound by CLSY3^40^. However, the role of this motif has not been investigated, leaving it unclear if this sequence is merely correlated with siren loci, which are themselves enriched near genes^41^ and thus are likely to overlap *cis*-regulatory elements, or if it is instructive in regulating the epigenome.

To identify additional genes required for RdDM at loci regulated by CLSY3, a forward genetic screen was designed to assess methylation levels at sites that lose methylation specifically in *clsy3* mutants. From this screen, and parallel reverse genetics approaches, we identified several REPRODUCTIVE MERISTEM (REM) family transcription factors (TFs) that are required for RdDM. In line with this new function in epigenetic regulation, we designate these REMs as REM INSTRUCTS METHYLATION (RIMs). While the RIMs all specifically affect loci regulated by CLSY3, mutations in *rim16* or *rim22* selectively affect HyperTE loci, which are targeted by RdDM in male reproductive tissue, while a *rim triple* CRISPR deletion (*rim-cr*; *rem11,12,46*) selectively affects siren loci, which are targeted by RdDM in female reproductive tissue. In total, these *rim* mutants affect ∼80% of HyperTE and ∼37% of siren loci, reducing siRNA levels to similar extents as observed in the *clsy3* mutant. These findings not only demonstrate the RIMs as major players in the RdDM pathway, but also reveal their sex-specific functions, making them unique compared to all other known RdDM factors. To investigate the mechanism linking the sequence-specific DNA binding activities of RIM transcription factors to epigenetic regulation, two orthogonal approaches were taken. On the female side, a DNA motif present at siren loci^40^ and bound by REM12^42^ was disrupted and found to be critical for CLSY3 recruitment and siRNA production. On the male side, key residues within the RIM22 DNA binding domain were mutated and shown to be critical for DNA methylation at HyperTE loci. These results link epigenetic regulation to genetic elements for the first time, uncovering a new mode of RdDM targeting and revealing how distinct patterns of methylation can be generated via mechanisms that fall outside the previously described self-reinforcing loops between chromatin modifications.

## Results

### RIM22 is a novel RdDM factor that specifically regulates loci controlled by CLSY3

To identify new factors required for RdDM at loci regulated by CLSY3 (*i.e.*, at *clsy3*-dependent loci) a forward genetic screen was conducted. This screen utilized a published collection of homozygous EMS mutants (HEM lines^43^) and a series of PCR-based methyl-cutting assays targeting either *clsy3*-dependent or *clsy1,2*-dependent loci in flower tissue (**Fig. 1A** and **Fig. S1A, B**). Low pass whole genome sequencing and allelism tests of several candidate mutants validated the utility of this screen by identifying new alleles of *nrpd1* and *nrpe1* that affected all loci tested, a new allele of *ago4* that specifically affected *clsy1,2*-dependent loci, and three new alleles of *CLSY3* that specifically affected *clsy3*-dependent loci (**Fig. S1C-E**). In addition to the mutations affecting known RdDM factors, a new factor was discovered. Though bearing no mutations in any of the known RdDM components^44^, ES1M5S10328 specifically affected *clsy3-*dependent loci. To identify this mutant, a mapping-by-sequencing approach was undertaken^45^. After backcrossing the ES1M5S10328 line to the un-mutagenized parent line from the HEM population, 13 F_2_ progeny that displayed methyl-cutting defects were pooled and sequenced, revealing a QTL on chromosome 3 as the causal region (**Fig. S1F**). This region was further refined by a larger F_2_ population containing 612 plants, resulting in a 1.03 Mb region containing 5 genes with missense mutations, including a proline to leucine mutation (Pro267Leu) within the putative DNA binding domain of an AP2/B3-like transcription factor, *REPRODUCTIVE MERISTEM 22 (REM22)*, hereafter designated as *REM INSTRUCTS METHYLATION 22* (*RIN22*) to reflect both its phylogeny and biological role (*rim22-3*; **Fig. S1F**). Methyl-cutting assays and an allelism test with a second allele of *RIM22* (SALK_091149^46^/*rim22-2*; **Fig. S1G**) confirmed a causal role for this gene in regulating DNA methylation at loci controlled by CLSY3 (**Fig. 1A**).

**Figure 1.**
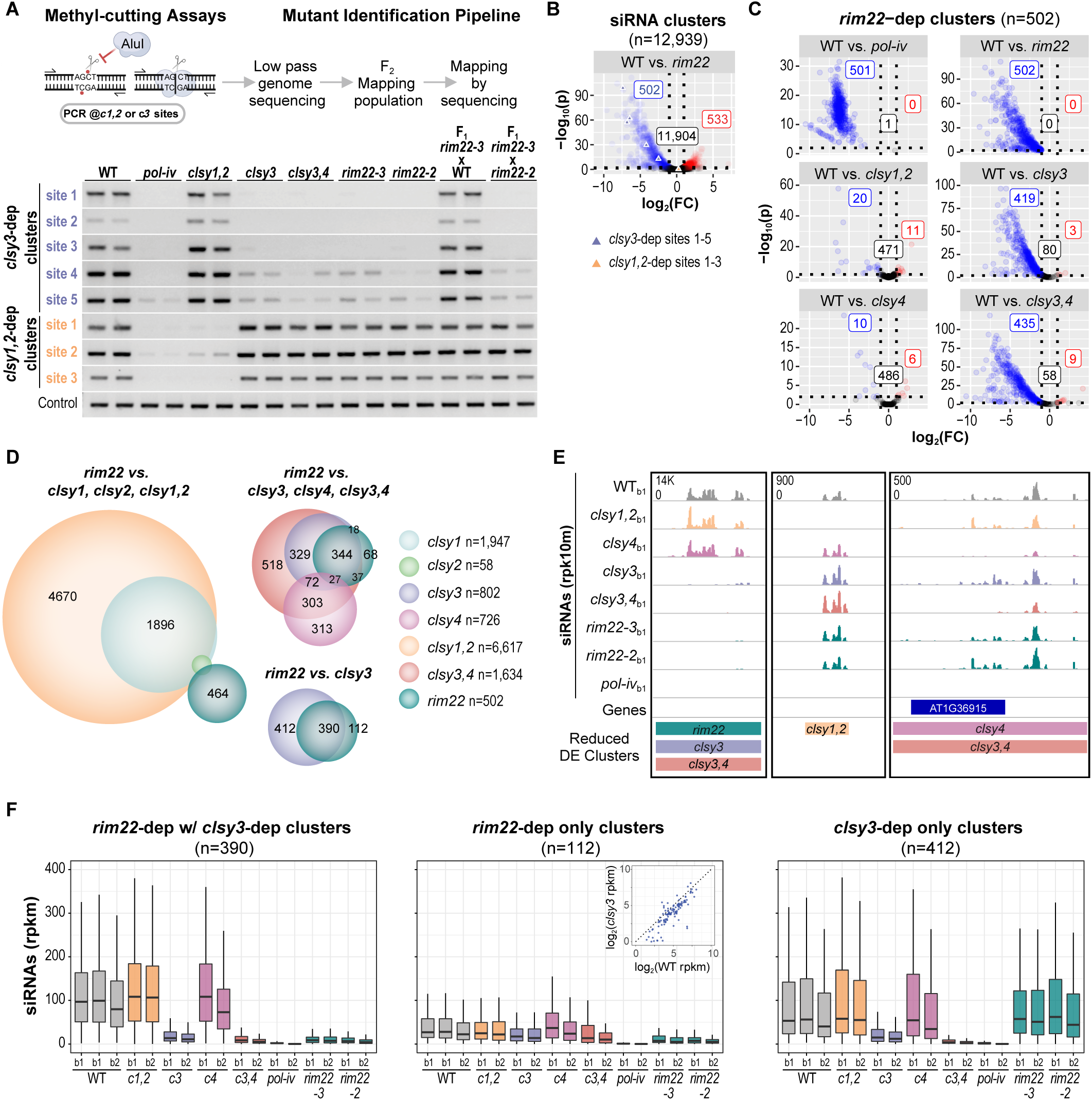
RIM22 is a new RdDM factor that is specifically required at a subset of *clsy3*-dependent loci. **(A)** (Upper) Cartoon schematic of a methyl-cutting assay to screen for DNA methylation mutants. The AluI restriction enzyme is inhibited by cytosine DNA methylation (red dots) within the AGCT motif allowing amplification by PCR across methylated motifs after AluI digestion. Alternatively, at unmethylated motifs, cleavage by AluI prevents amplification by PCR. Depending on the primers (half arrows) used for amplification, methylation can either be assessed at sites that lose methylation in *clsy1,2* mutants (*c1,2-*dep clusters) or *clsy3* mutants (*c3-*dep clusters). Mutants affecting methylation were sequenced and/or mapped to identify causal mutations. (Lower) Gels showing the amplification of DNA at the indicated sites after digestion with AluI. Two replicates are shown for the genotypes indicated above each set of lanes. The control shows the uniform amplification of a DNA region without the AluI restriction site. **(B and C)** Volcano plots showing siRNA levels at the full set of RdDM regulated clusters (n=12,939) defined in Zhou *et al.*^40^ **(B)** or just the 502 *rim22*-epenent clusters **(C)**. For each plot, clusters that are downregulated compared to wild-type (WT) controls (log_2_FC ≤ -1 and FDR < 0.01) are shown as blue circles, those unaffected are shown as black circles, and those upregulated (log_2_ FC ≥ 1 and FDR < 0.01) are shown as red circles. Samples used for the DEseq analysis are listed in **Supplementary Data 1.** (The number of clusters in each category are indicated in the correspondingly colored boxes and in **(B)** loci corresponding to the sites used for the methyl-cutting assay in **(A)** are highlighted by the different colored triangles. **(D)** Scaled Venn diagrams showing the relationships between reduced siRNA clusters in *rim22* and clusters previously shown to depend on different *clsy* mutants^40^. The circles are colored as indicated on the right and only overlaps >20 are labeled. **(E)** Screenshots showing siRNA levels at representative loci identified as being reduced in the indicated mutants from the differentially expressed cluster analyses (Reduced DE Clusters). For each locus, the siRNA expression range for all tracks is indicated in the upper left corner of the top track and the genotypes, as well as the smRNA-seq batch numbers (b1), are indicated on the left. **(F)** Boxplots showing siRNA levels at the indicated clusters. The genotypes and smRNA-seq batches (b1 or b2) are indicated below, with each batch representing a distinct biological replicate. Here, and for all subsequent boxplots, the graphs show the interquartile range (IQR), with the median shown as a black line and the whiskers corresponding to 1.5 times the IQR. Boxplot outliers were omitted to optimize visualization due to the wide range of data scales. A scatter plot is included as an inlay in the middle plot showing the expression levels (rpkm) of siRNAs at each of the 112 loci in the *clsy3* mutant compared to its wild-type control.

To determine how broadly RIM22 acts within the RdDM pathway, the levels of siRNAs in flower tissue were profiled genome-wide via small RNA-sequencing (smRNA-seq) experiments (**Supplementary Data 1**). Across a previously curated set of siRNA clusters regulated by RdDM (n=12,939)^40^ the effects of the two *rim22* mutants (*rim22-2* and *rim22-3*) were assessed together via batch corrected differential expression analyses relative to three wild-type (WT) controls. In the *rim22* mutants, siRNAs were significantly reduced at 502 clusters (including the 5 clusters overlapping the sites used for the methyl-cutting assay; blue triangles, **Fig. 1B** and **Supplementary Data 2**) and modestly increased at 533 clusters (**Fig. 1B** and **Supplementary Data 3**). These findings demonstrate an important role for RIM22 in regulating siRNA production at a subset of RdDM loci.

Comparing the effects of the *rim22* mutants on siRNA levels to those observed in *pol-iv* and various *clsy* mutants profiled in parallel demonstrated that, as expected, siRNA levels at the 502 *rim22*-dependent clusters were strongly reduced in a *pol-iv* mutant (**Fig. 1C**). However, they were differentially affected in the *clsy* mutants: While the vast majority of *rim22-*dependent clusters were also reduced in *clsy3* or *clsy3,4* mutants, very a few clusters were reduced in *clsy1,2* or *clsy4* mutants (**Fig. 1C**). This specificity for *clsy3*-dependent loci was also observed when the *rim22-*dependent clusters were overlapped with the full set of previously defined *clsy*-dependent categories from flower tissue^40^ (**Fig. 1D**): Very few *rim22*-dependent clusters overlapped with *clsy1-*, *clsy2-*, *clsy4*- or *clsy1,2*-dependent clusters (**Fig. 1D**), while ∼78-86% overlapped with *clsy3*- (390/502) and/or *clsy3,4*-dependent (434/502) clusters, respectively (**Fig. 1D** and **E**).

Having demonstrated the specificity of RIM22 for *clsy3*-dependent loci, the strength of the *rim22* mutants were assessed by quantifying siRNA levels over clusters co-regulated by *rim22* and *clsy3* (n=390), as well as those classified as only dependent on *rim22* (n=112) or *clsy3* (n=412). This analysis revealed that RIM22 has the strongest effect on loci that also depend strongly on CLSY3 and that this subset of CLSY3 targets are the most highly expressed group of clusters (**Fig. 1F**, n=390).

Furthermore, it demonstrated that even the loci classified as only dependent on *rim22* are also reduced in *clsy3* mutants (**Fig. 1F**, n=112 boxplot and scatter plot inlay)—the reductions were just too weak to pass the significance filter from the differential expression analysis. Lastly, it revealed that approximately half of *clsy3*-dependent loci in flower tissue are indeed fully independent of RIM22 (**Fig. 1F**, n=412). Together, these findings establish RIM22 as a critical new player in the RdDM pathway that acts specifically at *clsy3*-dependent loci and preferentially affects clusters that express the highest levels of siRNAs.

### RIM22 regulates *clsy3*-dependent RdDM targets that are highly expressed in male, but not female reproductive tissue

Amongst the *clsy3*-dependent clusters, distinct subsets are known to produce high levels of siRNAs in female and male reproductive tissues: siren clusters in ovules (n=133 as defined in Zhou *et al.*^40^) and HyperTE clusters in anthers (n=797 as defined in Long *et al.*^39^, equivalent to 715 of the 12,939 clusters from Zhou *et al.*^40^, **Supplementary Data 4**). Overlap analyses using these highly expressed clusters revealed that *rim22* affects siRNAs at ∼2/3rds of HyperTE loci (n=434/715), but only one siren locus (**Fig. 2A**). Consistent with previous characterization of HyperTE loci, siRNAs at these clusters were strongly reduced in *clsy3*, *clsy3,4* and *pol-iv* mutants, but largely unaffected in *clsy1,2* or *clsy4* mutants (**Fig. 2B**). In *rim22* mutants, strong reductions in siRNA levels were observed at HyperTE loci (**Fig. 2B**, Left), and these reductions are largely specific to the subset that overlap *rim22*-dependent clusters (n=434) (**Fig. 2B**, Middle). At these 434 HyperTE loci, *rim22* mutants mimic *clsy3,4* mutants, while at the non-overlapping HyperTE loci (n=281) siRNA levels were similar to the WT controls (**Fig. 2B**, Right). In agreement with this specificity, MethylC-seq experiments using flower tissue from *rim22* mutants (**Supplementary Data 5**) revealed that the strongest reductions in CHH methylation (n=320 DMRs; **Fig. 2C** and **Supplementary Data 6**) and CHG methylation (n=12 DMRs; **Fig. 2C** and **Supplementary Data 6)** essentially all overlap with HyperTE loci (**Fig. 2D**). At these DMRs, methylation levels are strongly reduced in the *rim22* mutants, approaching the levels observed in *pol-iv* mutants (CHH; **Fig. 2E** and CHG; **Fig. S2A**). To determine if more subtle changes in methylation that failed to pass the thresholds for DMR calling also occur in *rim22* mutants, the levels of CHH and CHG methylation were assessed across the different categories of siRNA clusters. Like for siRNAs, the *rim22* mutants displayed the strongest reductions in non-CG methylation, across the largest fraction of clusters (**Fig. 2F** scatterplots), at the 434 HyperTE loci that overlap *rim22*-dependent siRNA clusters (**Fig. 2F** and **Fig. S2B**). Much weaker effects were observed in *rim22* mutants at the remaining 281 HyperTE loci (**Fig. 2F** and **Fig. S2B**) and the patterns of methylation were similar to the wild-type controls at siren loci (**Fig. S2C**), *clsy1,2*-dependent clusters (**Fig. S2D**) and *clsy4*-dependent siRNA clusters (**Fig. S2E**). Together, these findings demonstrate that RIM22 activity is required for siRNA production and DNA methylation at HyperTE loci, which are actively targeted by RdDM in male reproductive tissues. However, RIM22 is not required at siren loci, which are targeted in female reproductive tissue. Taken together, these findings demonstrate that RIM22 is unique amongst RdDM factors as it is the only component of this pathway known to act in a sex-type specific manner.

**Figure 2.**
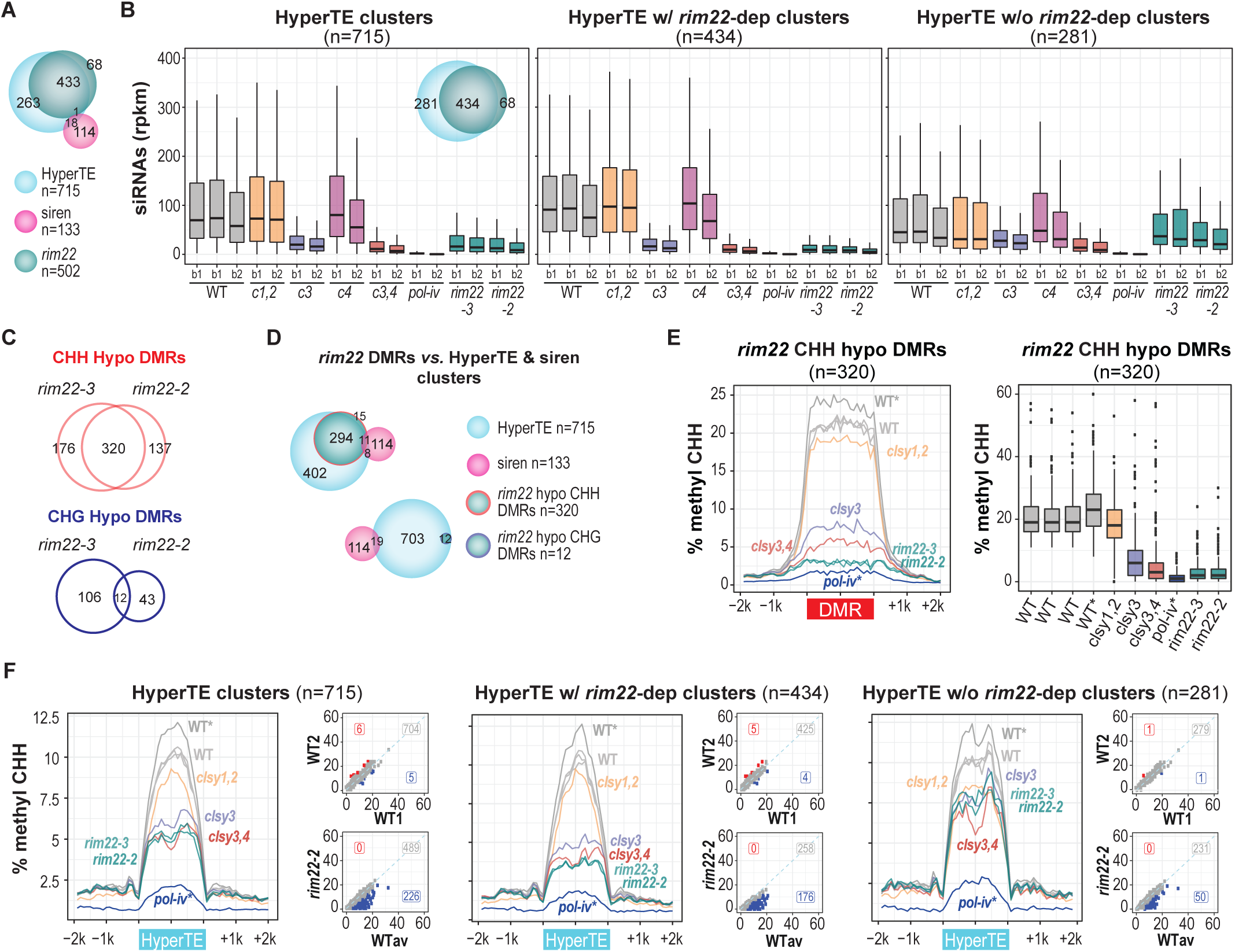
RIM22 is required for siRNA production and DNA methylation specifically at HyperTE loci. **(A)** Scaled Venn diagrams showing the relationships between siRNA clusters reduced in *rim22* and clusters previously shown to be highly expressed in ovules (siren loci)^40^ or anthers (HyperTE loci)^39^. The circles are colored as indicated below. **(B)** Boxplots showing siRNA levels at the indicated clusters. A scaled Venn diagram showing the different categories being compared is shown as an inlay in the first plot. The genotypes and smRNA-seq batches (b1 or b2) are indicated below, with each batch representing a distinct biological replicate. Boxplot outliers were omitted to optimize visualization due to the wide range of data scales. **(C)** Scaled Venn diagrams showing the overlap of DMRs (100 bp) identified in the *rim22-2* and *rim22-3* mutants for the CHH and CHG contexts. **(D)** Scaled Venn diagrams showing the overlap of shared *rim22* DMRs in the CHH and CHG contexts with HyperTE and siren siRNA clusters. The circles are colored as indicated on the right. **(E)** Metaplot (left) and boxplot (right) showing the levels of CHH methylation over the shared set of CHH DMRs identified in the *rim22* mutants. The metaplot shows the percent methylation in 100 bp bins across the DMRs and the flanking 2kb regions in the indicated genotypes. The boxplot shows the percent methylation at each DMR in the genotypes indicated below. For both plots, the two samples from a previous experiment (Zhou *et al.*^18^) are marked with asterisks (*). **(F)** Metaplots, as described in **(E),** showing the levels of CHH methylation over at the indicated sets of HyperTE loci. The scatter plots beside each metaplot show the changes in methylation between two representative WT samples (WT1 and WT2) or the average of 3 WT samples (WTav) and the indicated mutant. The number of clusters with higher (mutant – WT > 5%), lower (mutant – WT < -5%), or unchanged levels of methylation are indicated in red, blue, and grey, respectively.

### Distinct REM family members regulate *clsy3*-dependent RdDM targets in male and female reproductive tissues

Since RIM22 is part of a large family of related proteins^47^ and only affects a subset of *clsy3*-dependent targets, reverse genetics approaches were taken to identify additional proteins important for RdDM. On the male side, *REM16* (hereafter designated as *RIM16*) was selected as a candidate for the regulation of additional HyperTE loci based on its expression pattern in anthers^48^. On the female side, a group of three tandemly oriented *REM* genes (*REM46*, *REM11*, and *REM12*) were selected based on their expression patterns in ovules^48^ and on the enrichment of REM12 from DNA Affinity Purification and sequencing (DAP-seq) data^42^ at nearly all the previously identified CLSY3 ChIP-seq peaks^40^ (**Fig. 3A, B** and **Supplementary Data 7**). mRNA sequencing (mRNA-seq; **Supplementary Data 8**) in flower tissue confirmed the disruption of the *RIM16* gene in the *rim16-3* allele and disruption of all three tandemly oriented *REM* genes (*REM46*, *REM11*, and *REM12*) in a CRISPR engineered deletion line, hereafter referred to as *rim-cr* (**Fig. S3A, B**). Differential expression (DE) analyses using these mutants, as well as the *rim22* mutant, revealed many DE genes (**Supplementary Data 9**). However, none of these genes have known links to DNA methylation (**Fig. S3A, B** and **Supplementary Data 9**). Using these alleles, methyl-cutting assays were conducted at the 5 previously described *clsy3*-dependent sites, which all overlap with HyperTE loci, as well as an additional site overlapping a siren locus. In *rim16-3*, methylation was reduced at 4 of the 5 HyperTE loci, but not at the siren locus. In contrast, the *rim-cr* mutant displayed the opposite phenotype, reducing methylation at the siren locus, but none of the HyperTE loci (**Fig. 3C**). These findings demonstrate that different REM family TFs act in connection with CLSY3 to regulate DNA methylation at HyperTE versus siren loci.

**Figure 3.**
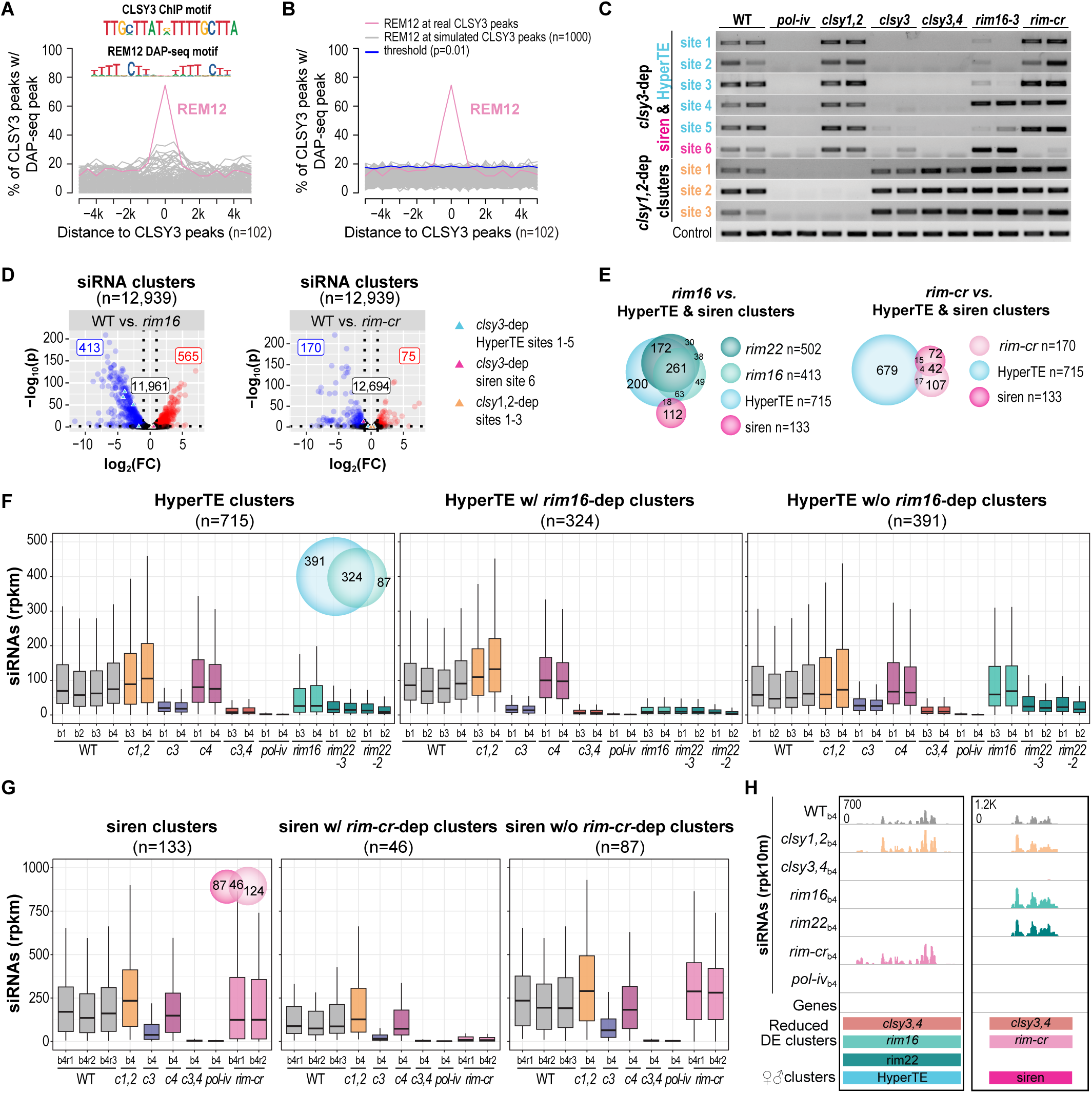
Distinct REM family TFs control RdDM at loci highly expressed in male versus female reproductive tissues. **(A)** Line plot showing the enrichment of DAP-seq peaks for the published set of Arabidopsis transcription factors^107^ at CLSY3 ChIP-seq peaks (n=102)^40^. Many of these peaks (n=66/102) contain one or more instances of the top motif identified from the CLSY3 ChIP-seq (CLSY3 ChIP motif) and most (n=94/102) correspond to siren loci^40^. The most enriched transcription factor, REM12, is labeled and highlighted in pink, while the other transcription factors are shown in grey. The REM12 DAP-seq and CLSY3 ChIP-seq motifs, which are quite similar, are shown above. **(B)** Line plot showing the enrichment of REM12 at real CLSY3 peaks or at a simulated set of CLSY3 peaks. This simulation was performed 1,000 times, and each one is shown as a grey line. The REM12 binding to the real CLSY3 peaks is highlighted in pink and the significance threshold (p=0.01) is shown in blue. **(C)** Methyl-cutting assay data for *rim16* and *rim-cr*. The gels show the amplification of DNA at the indicated sites after digestion with AluI, a methylation sensitive restriction enzyme. Two replicates are shown for the genotypes indicated above each set of lanes. The control shows the uniform amplification of a DNA region without the AluI restriction site. **(D)** Volcano plots showing siRNA levels at the full set of RdDM regulated clusters (n=12,939) defined in Zhou *et al.*^40^ in *rim16* and *rim-cr* mutants. For each plot clusters that are downregulated compared to wild-type (WT) controls (log_2_FC ≤ -1 and FDR < 0.01) are shown as blue circles, those unaffected are shown as black circles, and those upregulated (log_2_ FC ≥ 1 and FDR < 0.01) are shown as red circles. Samples used for the DEseq analysis are listed in **Supplementary Data 1.** The number of clusters in each category are indicated in the correspondingly colored boxes and loci corresponding to the sites used for the methyl-cutting assay are highlighted by the different colored triangles. **(E)** Scaled Venn diagrams showing the relationships between siRNA clusters reduced in *rim16*, *rim22,* or *rim-cr* and clusters previously shown to be highly expressed in ovules (siren loci)^40^ or anthers (HyperTE loci)^39^. The circles are colored as indicated on the right. For the Venn diagram on the right some overlaps are missing due to special constraints. **(F-G)** Boxplots showing siRNA levels at the indicated clusters. A scaled Venn diagram showing the different categories being compared is shown as an inlay in the first plot. The genotypes, smRNA-seq batches (b1, b2, b3, or b4), and biological replicates (r1, r2, or r3) are indicated below, with each batch representing a distinct biological replicate. Boxplot outliers were omitted to optimize visualization due to the wide range of data scales. **(H)** Screenshots showing siRNA levels at representative HyperTE and siren loci. For each locus, the siRNA expression range for all tracks is indicated in the upper left corner of the top track and the genotypes, as well as the smRNA-seq batch numbers (b4) are indicated on the left. Below each set of tracks the regions corresponding to the HyperTE and siren clusters are marked and the genotypes where these clusters were identified as being differentially expressed (Reduced DE Clusters) are indicated.

As was done for *rim22*, the effects of the *rim16* and *rim-cr* mutants on global siRNA levels were determined via smRNA-seq experiments using flower tissue and compared to various combinations of *clsy* mutants. Across the previously curated set of siRNA clusters regulated by RdDM (n=12,939^40^), *rim16* mutants showed 413 reduced clusters (including 4 of the 5 loci used in the methyl-cutting assay) and 565 increased siRNA clusters (**Fig. 3D**). For the *rim-cr* triple mutant, 170 reduced clusters (including the siren locus used in the methyl-cutting assay) and 75 increased siRNA clusters were identified (**Fig. 3D**). As expected, siRNA levels at the 413 *rim16*- and 170 *rim-cr*-dependent clusters were strongly reduced in a *pol-iv* mutant (**Fig. S3C, D**). As observed for *rim22*, the effects of *rim16* and *rim-cr* mutants were specific to *clsy3*-dependent loci (**Fig. S3C-F**) and displayed the strongest effects at highly expressed, *clsy3*-dependent loci (n=300 or n=103, respectively; **Fig. S3G, H**). Consistent with these findings, and the results from the methyl-cutting assays, the *rim16*-dependent clusters overlapped with HyperTE clusters (and *rim22*-dependent clusters), but not with siren loci. In contrast, *rim-cr* displayed the opposite behavior, overlapping with about 1/3^rd^ of the siren loci but with very few HyperTE clusters (**Fig. 3E**). Attesting to the crucial roles and high degree of locus specificity for these RIMs, the *rim16* and *rim-cr* mutants mimicked the strong reductions in siRNA levels observed in *clsy3* and *clsy3,4* mutants at specific subsets of HyperTE (n=324) and siren (n=46) loci, respectively (**Fig. 3F, G**) but showed minimal effects outside these loci (**Fig. 3F-H** and **Fig. S3 I, J**). Together, these findings demonstrate that distinct REM family TFs are critical for regulating siRNAs and DNA methylation at *clsy3*-dependent loci that are specific to male and female reproductive tissues.

### Additional factors are required for siRNA production at some siren and HyperTE loci

Although many HyperTE and siren loci are strongly dependent on either RIM16, RIM22 or the RIM-CR triple mutant, respectively, others remain expressed at wild-type levels (**Figs. 2B, 3F** and **3G**), suggesting either more factors are required, or these RIMs act redundantly to control RdDM at the remaining HyperTE and siren loci. To distinguish between these possibilities, higher order combinations of REM mutants were generated. Like the *rim* single mutants, flower smRNA-seq experiments identified both up and downregulated siRNA clusters in the *rim16,22* double and *rim* quintuple (*rim16,rim22,rim-cr*) mutants (**Fig. S4A**). Comparing the reduced siRNA clusters in the *rim16* and *rim22* single mutants with the *rim16,22* double mutant revealed a complex genetic relationship between these TFs in regulating HyperTE, but not siren loci (**Fig. 4A**). The largest category of HyperTE loci (n=261) were strongly reduced in either the *rim16* or *rim22* single mutants, with no clear synergistic effect in the *rim16,22* double (**Fig. 4A, B** and **Fig. S4B**), demonstrating that both these TFs are critical for siRNA production at these loci. However, some loci are predominantly reduced in *rim22* but not in *rim16* (n=173) or vice versa (n=63) (**Fig. 4A, B** and **Fig. S4B**), revealing these TFs can also act independently. Finally, while these TFs do act synergistically at some loci, including 71 HyperTE loci, many HyperTE loci remain unaffected in the *rim16,22* double mutant (n=147) (**Fig. 4A, B** and **Fig. S4B**). Indeed, these loci also remain unaffected even in a *rim* quintuple mutant, confirming that additional factors are required to regulate siRNA production at some HyperTE loci. Interestingly, many of the HyperTE loci in this category are reduced in *clsy1,2* but not *clsy3,4* mutants, suggesting multiple factors are missing, some, like REM16 and REM22, that act with CLSY3 and others that act with CLSY1 and/or CLSY2 to control siRNA production at HyperTE loci.

**Figure 4.**
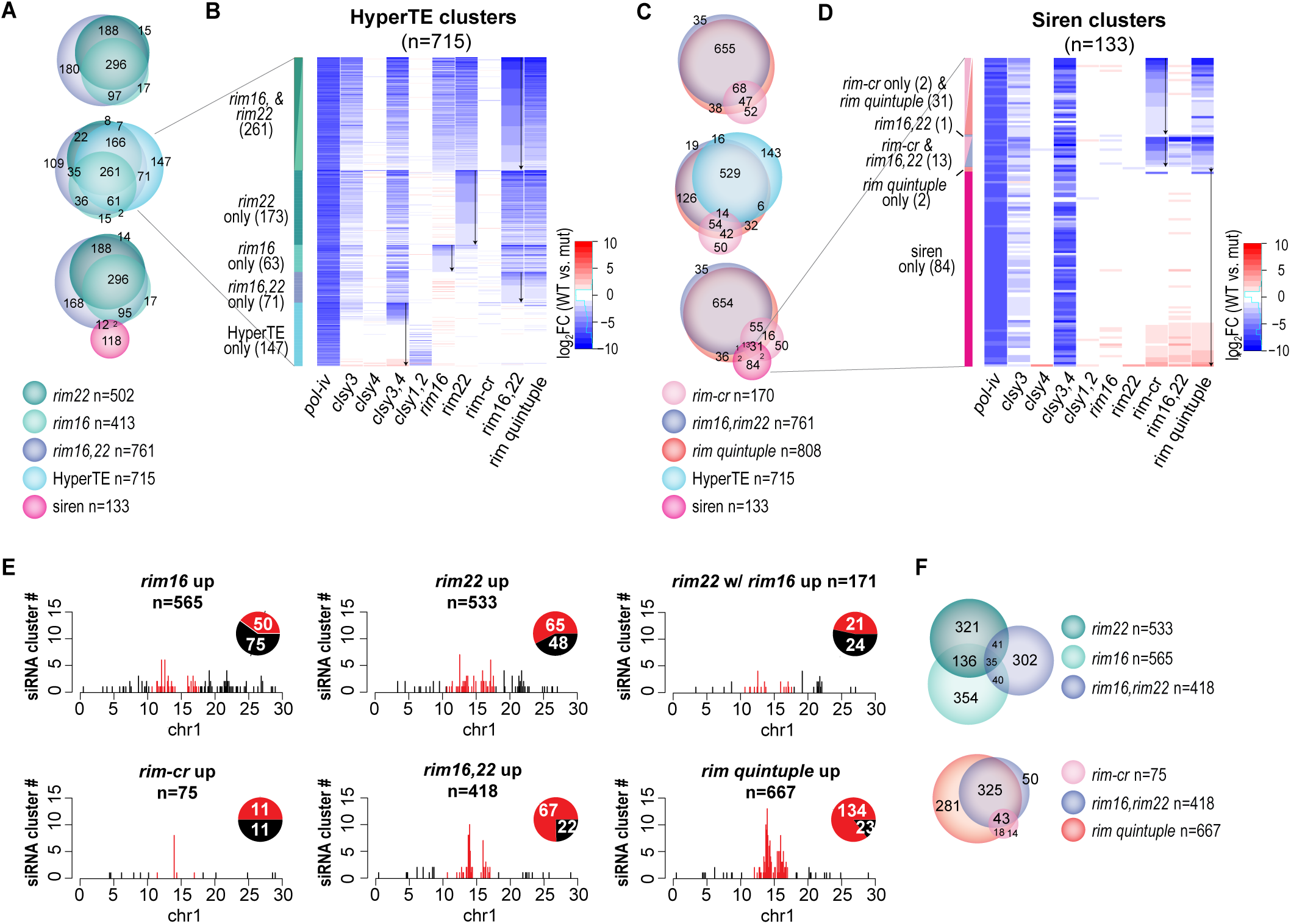
Limited redundancy is observed between RIM TFs at HyperTE and siren loci but higher order mutants do enhance siRNA production in pericentromeric heterochromatin. **(A** and **C)** Scaled Venn diagrams showing the relationships between reduced siRNA clusters in the indicated *rim* mutants and either HyperTE or siren loci. The circles are colored as indicated below. Some minor overlaps (n<5) are missing due to special constraints. **(B** and **D**) Heatmaps showing the log_2_FC in siRNA levels between WT controls and the mutants indicated below each column. The heatmaps are grouped based on the scaled Venn diagrams including HyperTE or siren loci from **(A)** and **(C)**, respectively. Each group is color-coded and labeled based on the mutants showing reduced siRNA levels. For each group, the clusters are in the same order for all genotypes and the order was determined based on a ranking (from most to least affected) of the genotypes marked with downward arrows. **(E)** Bar and pie charts showing the distributions of upregulated siRNA clusters in the indicated genotypes across chromosome 1 (chr1). The distributions across the other chromosomes are shown in **Fig. S5**. The total number of clusters across all five chromosomes is indicated above each plot. For both the bar and pie charts, the red and black colors indicate pericentromeric heterochromatin and chromosome arms, respectively. The numbers in the pie charts represent the number of clusters in the corresponding category for chr1. **(F)** Scaled Venn diagrams showing the relationships between upregulated siRNA clusters in the indicated *rim* mutants. The circles are colored as indicated on the right.

Like the behavior at HyperTE loci, limited redundancy was observed between the *rim16* and *rim22* single mutants and the *rim-cr* triple mutant at siren loci (**Fig. 4C, D** and **Fig. S4C**). A few siren loci were reduced in the *rim16,22* double, but most were weaker than observed for *rim-cr* triple, and nearly 2/3rds of siren loci (n=84) remain expressed at or above wild-type levels in the *rim* quintuple mutant (**Fig. 4C, D** and **Fig. S4C**). Taken together, these findings confirm the specificities of RIM16 and RIM22 for the regulation of HyperTE loci and RIM-CR *(i.e.*, REM11/REM12/REM46) for the regulation of siren loci, which is consistent with their distinct expression patterns in male and female reproductive tissues, respectively. These data also signal the existence of additional factors that are required to regulate the remaining HyperTE (n=147/715) and siren (n=83/133) loci, suggesting a complex network of factors are needed to regulate these reproductive-specific siRNA loci.

### RIM TFs are required to prevent hyper-siRNA production on a global scale

In addition to reduced siRNA expression at HyperTE and siren loci, the *rim* mutants show enhanced siRNA production at hundreds of RdDM targets (**Figs. 1B, 3D, S4A,** and **Supplementary Data 3**). As evidenced by the heatmaps in **Fig. 4B** and **4D**, very few of these upregulated siRNA clusters correspond to HyperTE or siren loci. For the *rim16, rim22, and rim-cr* mutants, the upregulated clusters are spread relatively evenly throughout the genome (**Fig. 4E and Fig. S5A**). Interestingly, the *rim16* and *rim22* upregulated siRNA clusters overlap to a lower extent than observed for their respective downregulated siRNA clusters (compare **Fig. 4A** and **Fig. 4F**) and this upregulation is suppressed in the *rim16,22* double mutant (**Fig. 4F**). Instead of a uniform distribution of upregulated clusters, the *rim16,22* double almost exclusively affects clusters within pericentromeric heterochromatin, a phenotype that is enhanced in the *rim* quintuple mutant (**Fig. 4E, F**). While the mechanism(s) underlying the observed over-production of siRNAs in *rim* mutants is unknown, these findings reveal roles for RIM TFs in promoting homeostasis within RdDM to ensure high levels of siRNAs are produced at HyperTE and siren loci and that other loci, especially those in pericentromeric heterochromatin, remain lowly expressed.

### Epigenetic regulation by the RIMs relies on genetic information, linking epigenetic targeting to DNA motifs

Our previous identification of a DNA motif (*i.e.,* the CLSY3 motif) present at many CLSY3 ChIP-seq peaks^40^, together with the identification of distinct TFs required for siRNA production at HyperTE and siren loci, suggests genetic information is critical for the regulation of loci controlled by CLSY3 in reproductive tissues. At the subset of CLSY3 peaks containing the CLSY3 motif (n= 66/133, 98.5% of which overlap siren loci^40^), REM12 is highly enriched over the motif, however, the levels of CLSY3, siRNAs, and CHH methylation are offset from the motif, instead showing enrichment at either up- or down-stream regions (**Fig. 5A** and **Supplementary Data 10**). Based on these observations, the CLSY3 motif could facilitate the recruitment of CLSY3-Pol-IV complexes to transcribe siRNAs from adjacent regions or it could act as a barricade to restrict the activity of RdDM and prevent the spreading of methylation. To test these hypotheses, siRNAs were profiled in lines with disruptions in four different CLSY3 motifs. In two lines T-DNA insertions were identified either within or adjacent to CLSY3 motifs (CS828917 and Salk_107926, respectively) and in two other lines the CLSY3 motifs were deleted using CRISPR (**Fig. 5B** and **Supplementary Data 1-3**). Notably, while REM12 binds all these loci *in vitro* (**Fig. 5A**), only 2 of the 4 show reduced siRNAs in the *rim-cr* mutant (**Fig. 5B**), suggesting additional genetic or epigenetic features shape the function of REM12 *in vivo*. Nonetheless, in all cases disruption of the CLSY3 motif resulted in drastic reductions in siRNA levels (5-100x reduced) at the targeted locus, mimicking the losses observed in *clsy3* or *clsy3,4* mutants (**Fig. 5B**). As minimal changes in siRNAs at other loci were observed (**Fig. S6A, B**), these findings are consistent with a critical, *cis*-regulatory function for the CLSY3 DNA motif. Furthermore, the low levels of siRNAs that remained at these disrupted loci showed similar patterns as observed in the wild-type samples (**Fig. 5B**), supporting a role in the recruitment of CLSY3-Pol-IV complexes rather than a role in blocking the spreading of RdDM. To further support this hypothesis, CLSY3 enrichment at one of the CRISPR-deleted motifs (peak 3-396) was assessed by ChIP-qPCR. (This CRISPR deletion was created in the same CLSY3-3xFLAG/*clsy3* transgenic line previously used in ChIP-seq experiments^40^ and shown to complement the *clsy3* mutation^40^). While CLSY3 enrichment was observed at this locus in the unedited background, it was lost in the CRISPR-deletion background (**Fig. 5C**). As a positive control, CLSY3 enrichment at another CLSY3 target (peak 4-336) was tested and found to be similarly enriched in both the edited and unedited backgrounds (**Fig. 5C**). These findings demonstrate a critical role for the CLSY3 DNA motif in the production of siRNAs and the recruitment of CLSY3 at siren loci, linking, for the first time, a genetic element to epigenetic regulation via the RdDM pathway.

**Figure 5.**
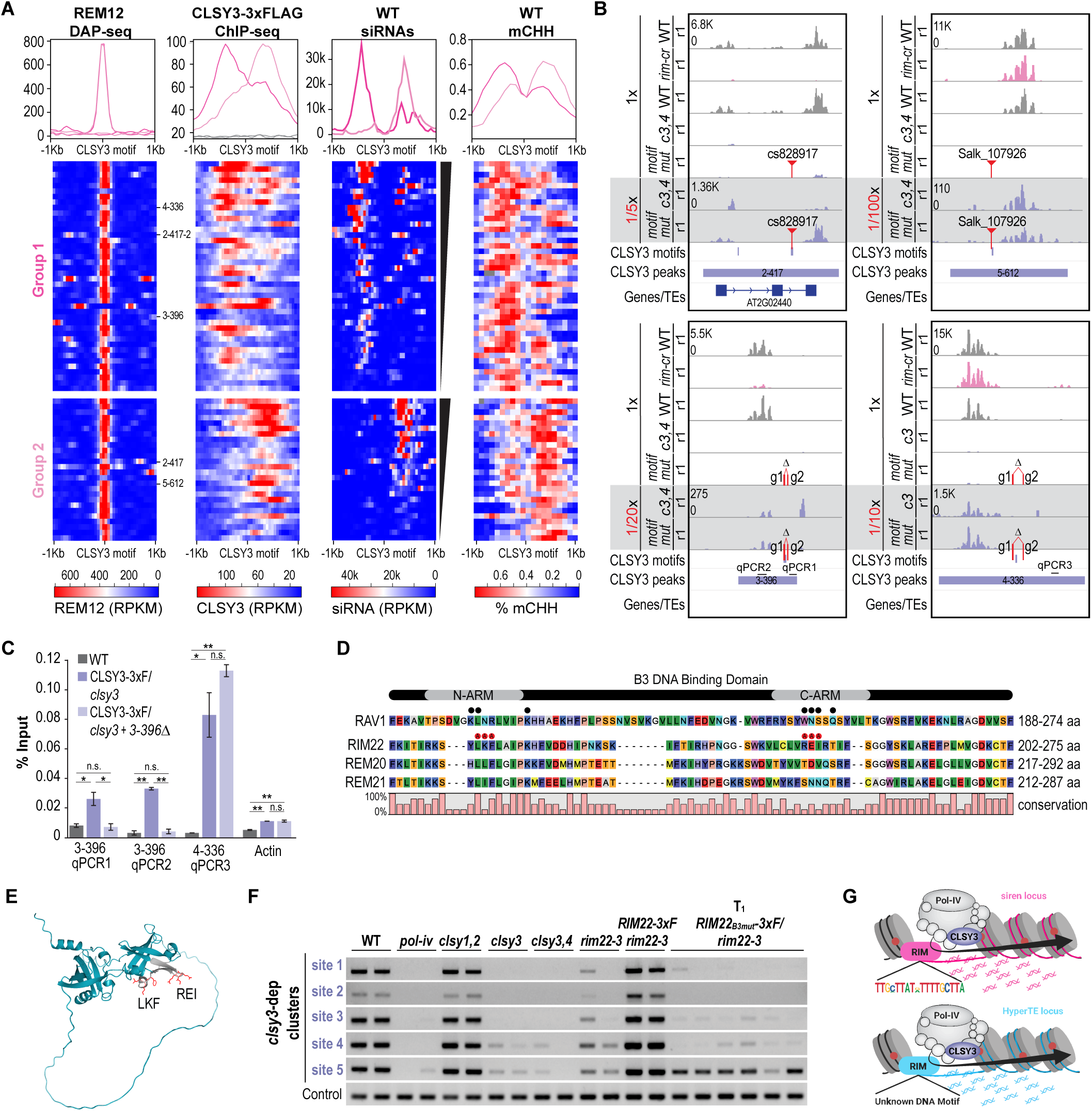
Epigenetic regulation by the RIMs relies on genetic information, linking epigenetic targeting to DNA motifs. **(A)** Profile plots and heatmaps showing the levels of REM12 enrichment, CLSY3 enrichment, siRNA levels, or CHH methylation (mCHH) in the 1 Kb regions surrounding CLSY3 motifs present under CLSY3 ChIP peaks. The scales for each set of plots are shown below. The plots are split into two groups and ranked (black triangles) based on the patterns of siRNAs expression such that all the plots are in the same order. For the REM12 DAP-seq and CLSY3 ChIP-seq, the profile plots are shown for the empty vector and WT controls (grey lines), respectively, but the heatmaps were omitted as no enrichment was observed. The profile plots for group 1 and group 2 are shown in pink and light pink, respectively. The CLSY3 peak identification numbers for the loci shown in **B** are marked. **(B)** Screenshots showing siRNA levels at siren clusters with disrupted CLSY3 motifs. For each locus, the positions of CLSY3 motif(s), CLSY3 peaks, and genes/TEs are shown below. In addition, the positions of the T-DNAs or gRNAs used to generate CRISPR deletions are shown in red. The tracks show siRNA levels on two scales for the indicated genotypes. A 1x scale to compare siRNA levels to the WT control and a variable, reduced scale (1/5x to 1/100x; grey background) to visualize the patterns of residual siRNAs in the motif mutant lines. **(C)** ChIP-qPCR showing CLSY3 binding to chromatin is blocked specifically at loci where the CLSY3 motif is disrupted. The error bars represents the standard deviation of two technique replicates. For each region tested (3-396 qPCR1, 3-396 qPCR2, and 4-336 qPCR3), the primer positions were shown in **(B).** Statistical significance was determined by Student’s *t*-test (* and ** represent *p*<0.05 and *p*<0.01, respectively). **(D)** Alignment of the B3 domains, including the N-ARM and C-ARM regions, from the indicated TFs. For each TF, the amino acids (aa) shown are indicated on the right and the conservation (0-100%) is shown below. For RAV1, the experimentally determined DNA interacting residues from Yamasaki *et al.*^51^ are marked with black triangles. For RIM22, the residues mutated to alanines are marked with red circles. – represents a gap in the alignment. **(E)** Alphafold prediction of RIM22^49,50^ (teal) with the N- and C-ARMs of the second B3 DNA binding domain shown in grey and the mutated residues within these regions shown in red. **(F)** Methyl-cutting assay data for two replicates of the indicated controls and six independent T_1_ lines expressing a mutated version of RIM22 (RIM22_B3mut_-3xF) introduced into the *rim22-3* background. The gels show the amplification of DNA at the indicated sites after digestion with AluI, a methylation sensitive restriction enzyme. The control shows the uniform amplification of a DNA region without the restriction site. **(G)** Model for RdDM at siren loci (upper) or HyperTE loci (lower) where targeting requires DNA sequences rather than chromatin modifications. Cartoons showing the binding of RIM TFs to the CLSY3 sequence motif (upper) or an unknown motif (lower), resulting in the recruitment of CLSY3-Pol-IV complexes to chromatin. Once targeted, the CLSY3-Pol-IV complexes initiate the production of siRNAs, resulting in the targeting of DNA methylation (red circles).

A sequence motif has not been identified for HyperTE loci. Thus, the importance of genetically encoded information in regulating these loci was tested by mutating the RIM22 B3 DNA binding domain. Based on the AlphaFold prediction of RIM22^49,50^ and its homology with RAV1, a B3 TF with known DNA-interacting residues^51^, B3 mutations were introduced into the predicted DNA-interaction loops within the N-ARM (*i.e.* LKF to AAA) and C-ARM (*i.e.* REI to AAA) of a *pRIM22::RIM22-3xFLAG* (*RIM22_B3mut_-3xF*) construct (**Fig. 5D, E**). For several independent T_1_ lines, methyl-cutting assays demonstrated that the *RIM22_B3mut_-3xF* construct failed to complement the *rim22-3* DNA methylation defects (**Fig. 5F**). Since the wild-type *pRIM22::RIM22-3xFLAG* (*RIM22-3xF*) construct complements the *rim22-3* mutant well (**Fig. 5F** and **Fig. S6B**), these findings demonstrate the importance of the RIM22 DNA binding domain for RdDM at HyperTE loci. They also support the notion that yet to be defined genetic information is required to instruct CLSY3 function at HyperTE loci, linking genetic elements to epigenetic regulation at loci regulated by CLSY3 during both male and female reproductive development.

## Discussion

From a forward genetic screen we uncovered a new mode of DNA methylation targeting that relies on several RIM TFs and their associated DNA motifs. In addition, we found this mode of targeting is restricted to loci that are regulated by CLSY3 and that different RIMs are required for RdDM at HyperTE loci, which are targeted in anthers, and siren loci, which are targeted in ovules. Both these findings transform our view of how DNA methylation patterns are regulated. First, as detailed in the introduction, all the previously identified mechanisms for regulating DNA methylation patterns—whether it be through the RdDM, CMT2/3, MET1, or demethylation pathways—rely on epigenetic rather than genetic features. Thus, our finding that DNA motifs and TFs are required to instruct methylation patterns represents a paradigm shift. Second, these findings reveal RIM TFs as the first sex-type specific regulators within the RdDM pathway, adding an additional layer of specificity, beyond the expression patterns of the CLSYs, that enables the generation of distinct methylation during plant development. Furthermore, our findings demonstrate that this new mode of DNA methylation targeting is a widespread mechanism since we have shown RIMs regulate approximately 80% of HyperTE loci and 37% of siren loci. Together these major advances establish a direct link between genetic elements and tissue-specific epigenetic regulation and, as discussed below, raise key questions, including how these connections are mediated at chromatin and whether additional REM family TFs are required to regulate methylation via the RdDM pathway.

In plants there are five families of B3 DNA binding domain TFs, and the REM family is the largest, but least well characterized^47,52^. Early studies demonstrated the REMs are expressed primarily in reproductive tissues, hinting at their functions^53–56^. However functional characterization has remained limited, at least in part due to their organization within the genome in multi-gene clusters^47,52^. Such organization suggests that REM genes may act redundantly, but unfortunately, this also makes them less genetically tractable. Thus far, a combination of genetic mutants, overexpression lines, and RNAi approaches have linked specific REMs to flowering time (REM5/VRN1^57^, REM13^58^, REM16/RIM16^59^, REM17/TFS1^60^, REM34^58^, and REM46^58^) and reproductive development (REM11/VAL^61^, REM20/VDD^62^, REM22/RIM22^63^, REM34^64^, and REM35^64^). Our work adds a third category: the regulation of DNA methylation patterns by REM16/RIM16, REM22/RIM22 and at least one of the REMs (REM11, REM12, or REM46) deleted in the *rim-cr* deletion line.

The overlap in REMs linked to flowering time, reproductive development, and RdDM is notable, suggesting dual roles in distinct biological processes for many REMs, including REM16/RIM16, REM22/RIM22, and possibly also REM11. In this respect, using Arabidopsis as a model system offers a unique opportunity to study the functions of REM factors in RdDM separate from their roles in reproductive development. This is because even the strongest RdDM mutants in Arabidopsis have normal fertility whereas similar mutants in many other plants (*i.e.* brassica^65^, maize^66,67^, rice^68–70^, and tomato^71^) display strong fertility defects. In the case of REM16/RIM16 and REM22/RIM22, we found their respective single mutants displayed strong epigenetic defects, mimicking the siRNA and DNA methylation losses observed in *clsy3* mutants, demonstrating non-redundant roles in regulating RdDM. However, these same alleles did not show defects in flowering time^59^ or reproductive development^53,56^. Rather, a REM16/RIM16 RNAi line that reduced the levels of several related REMs^59^ and the *rem22-1/rim22-1* allele, where a T-DNA in the *REM22/RIM22* promoter caused an overexpression phenotype^63^, were required to affect flowering time^59^ and ovule development^63^, respectively. Thus, not only can REMs act in different processes, but their genetic relationships within these roles can also differ such that for some processes they act independently, while in others they play redundant roles. In the case of REM11, it remains to be determined whether this REM alone, or in combination with REM12 and/or REM46 regulates DNA methylation patterns. Nonetheless, as our *rim-cr* mutant failed to display the ovule fertilization defects observed in REM11 RNAi lines^61^, our findings suggest REM11 may be acting redundantly with other closely related REMs to control fertilization. In moving forward, it will be important to determine if any of the many other REMs regulate methylation at HyperTE or siren loci in connection with CLSY3, or if they can instead be linked to loci regulated by other CLSY family members. Furthermore, it will be interesting to see whether it remains a trend that these factors have dual roles in regulating plant development and controlling DNA methylation patterns.

Beyond showing RIM TFs are important for RdDM, our data demonstrates that key residues within their B3 DNA binding domains and the DNA motifs they recognize are critical for regulating RdDM in plant reproductive tissues—marking the first direct links between genetic elements and the targeting of DNA methylation. Furthermore, our motif deletion assays demonstrate that these DNA elements act in *cis*, regulating CLSY3 recruitment and siRNA production at the associated locus, but not other RdDM loci. Based on these findings, and our observation that the binding profile of REM12, and its associated DNA motif, are offset from the production of siRNAs and the accumulation of DNA methylation at siren loci, we propose the following non-exclusive working model (**Fig. 5G**). RIM TFs are recruited to RdDM loci via the preferences of their B3 DNA binding domains where they either interact transiently with CLSY3-Pol-

IV complexes (CLSY3^72^ and Pol-IV^73^ affinity purification and mass spectrometry experiments show no stable interactions with RIM TFs) or they modify chromatin in a manner required for the subsequent recruitment of CLSY3-Pol-IV complexes. Once targeted, the CLSY3-Pol-IV complexes act in a directional manner to produce DNA methylation-targeting siRNAs either up or downstream of the RIM binding sites.

While the data behind our models provides significant insights into the mechanisms linking RIM TFs to epigenetic regulation via the RdDM pathway, key questions remain at many steps of the process. For example, the B3 DNA binding domains in the REM family of TFs have been proposed to act in both sequence specific and non-sequence specific manners and these factors have also been shown to act alone or as heterodimers^47,52^, raising questions about how and in what combinations different TFs are recruited to chromatin at different sets of RdDM targets. Attesting to the importance of these considerations, comparisons of the DAP-seq binding profile of REM12 with our siRNA profiling in the *rim-cr* triple, demonstrate that REM12 does not control siRNA production *in vivo* at all loci it binds *in vitro*. This discrepancy could indicate that REM12 binding is affected by the chromatin state in which its motif resides or that REM12 acts redundantly with other REMs at some loci, but not others, possibly depending on the presence or absence of other DNA motifs. After binding to chromatin, questions remain about how the RIM TFs coordinate the recruitment of CLSY3-Pol-IV complexes. Given the directionality observed for siRNA production, we favor the hypothesis that the RIMs transiently interact with CLSY3-Pol-IV complexes, acting akin to a Pol-II initiation complex, to orient Pol-IV and allow unidirectional transcription. However, it remains possible that the RIMs act indirectly by modifying chromatin and/or recruiting other factors that promote recruitment and directional transcription of CLSY3-Pol-IV complexes (**Fig. 5G**). Addressing these questions for REM12, as well as the other RIMs linked to RdDM, will be a critical next step that will shed light on the rules and cis-regulatory syntaxes decoded by the RIMs to link genetic elements to tissue-specific epigenetic regulation via the RdDM pathway.

## Acknowledgments

We thank colleagues and lab members for their comments on the manuscript. J.A.L. was supported by the NIH (GM112966). G.X. was supported by a postdoctoral fellowship from the Paul F. Glenn Center for Biology of Aging Research at the Salk Institute. F.W. was supported by a Pioneer postdoctoral fellowship from the Salk Institute. This work was also supported by the NGS Core Facility of the Salk Institute with funding from NIH-NCI CCSG: P30 CA01495, NIH-NlA San Diego Nathan Shock Center P30 AG068635, the Chapman Foundation and the Helmsley Charitable Trust.

## Author contributions

J.A.L. and G.X. designed the research. G.X. and Y.C. generated the genetic materials. G.X conducted experiments. G.X. and J.A.L analyzed and interpretated the data. F.W., and E.L. developed part of the methyl-cutting assays. J.A.L and G.X wrote the manuscript. All authors read and commented on the manuscript.

## Data availability

Illumina sequencing of smRNA-seq, mRNA-seq, and methylC-seq datasets generated in this study were deposited in the NCBI Gene Expression Omnibus (GEO) and are accessible via the accession number GSE###. Published data sets used in this manuscript include: smRNA-seq and CLSY3 ChIP-seq data from GSE165001^40^ and REM12 DAP-seq data from GSE60143^42^.

## Materials and methods

### Plant materials and growth conditions

All materials used in this study were in the Columbia-0 (Col-0) background. Previously published mutant lines include: *nrpd1-4* (SALK_083051)^11^, *nrpe1-12* (SALK_033852)^74^, *ago4-5* (EMS mutant)^75^, *clsy1-7* (SALK_018319)^76^, *clsy2-1* (GK_554E02)^18^, *clsy3-1* (SALK_040366)^18^, and *clsy4-1* (SALK_003876)^18^. The *rim22-2* (SALK_091149)^46^ and *rim16-3* (WiscDsLox443A7)^77^ mutants, as well as the two lines with insertions in CLSY3 motifs (SALK_107926^46^ and (CS828917/SAIL_663_H02^78^), were ordered from the Arabidopsis Biological Resource Center (ABRC). The deletion line disrupting the *REM11,REM12,REM46* cluster was generated using CRISPR (see **CRISPR/Cas9 editing** for details). The HEM EMS-mutagenized population was developed and characterized by Capilla-Perez *et al.*^79^ and Carrere *et al.*^44^.

For transgenic lines that required drug selection, seeds were sterilized in sterilization buffer (70% ethanol, 0.05% SDS) for 10 minutes, followed by plating on half strength Linsmaier and Skoog medium (1/2 LS; Caisson Labs, Cat# LSP03) supplemented with 0.6% agar (Phyto Agar, Cat# P1003). The plates were placed in a 4_°_C cold room for 4 days before growing in a growth chamber with 16h-light and 8h-dark cycles for 7 days. The seedlings were transplanted and grown under Salk greenhouse conditions. For HEM EMS lines, T-DNA insertion lines, and their corresponding control lines, seeds were stratified in 0.1% agar (Phyto Agar, Cat# P1003) at 4_°_C for 4 days before sowing directly onto soil and grown under Salk greenhouse conditions. Approximately five weeks after planting, stage 12 and younger flower buds were collected and frozen in liquid nitrogen.

### CRISPR/Cas9 editing

The *REM11/REM12/REM46* gene cluster was deleted (*rim-cr*) using a two-gRNA CRISPR/Cas9 gene editing system^80^. The gRNA design was performed using the CRISPR-PLANT online tool (http://omap.org/crispr/CRISPRsearch.html). The spacers (AGATGGGTAATTCCTGAAG and TGAAGGACGAAGATCTCAA) and the gRNA scaffold were PCR amplified from the pCBCdT1T2 plasmid^81^ and cloned into the BsaI site of the pHEE401E^80^ vector, which contains an egg cell driven zCas9 gene using Golden Gate cloning per manufacturer’s instructions (NEB, Cat# E1601). The constructs were transformed into AGLO agrobacteria competent cells and transformed into Arabidopsis Col-0 plants using the floral dip method^82^. Positive T_1_ seeds were selected under a microscope (ZEISS, AXIO Zoom.V16) equipped with a FS 38 HE filter to identify seeds expressing mCherry. Deletions were identified in the T_2_ generation using the PCR primers listed in **Supplementary Data 11**. As the two gRNAs used to generate the deletion are located within an intron of REM12 and an exon of REM46, respectively, the resulting cDNA from the *rim-cr* homozygous mutant flowers were sequenced to confirm that the deletion resulted in a nonfunctional transcript. Briefly, total RNA was extracted from flower buds of the CRISPR deletion line using the ZYMO RNA mini prep kit (Zymo Research Corporation, Cat# 50-444-597). The cDNA was obtained by reverse transcription using the Applied Biosystems High-Capacity cDNA Reverse Transcription Kit (Applied Biosystems, Cat# 4368814) and used to amplify the fusion transcript using the primers listed in **Supplementary Data 11**, which flank the two gRNAs. The PCR product was cloned into Blunt II TOPO vector (Invitrogen, Cat# 451245), transformed into TOP10 *E. coli* competent cells (Invitrogen, Cat# C404010), and three clones were subject to plasmid mini prep (BioPioneer Inc, Cat# PMIP-100) and sequenced at Primordium Labs. The sequences were aligned using SnapGene (v7.0.2).

The motifs under CLSY3 peaks 4-336 and 3-396 were deleted using the same two-gRNA CRISPR/Cas9 system and gRNA design tool described above. For the motifs under the 4-336 and 3-396 peaks the spacer sequences used were (CGATAAAACATTTAAATGAG and CGGAGACGAAACTATATCTT) or (AGCAAAATAGAAGCAAATAT and TATGTTTCTTGATCTAATTT), respectively. The agrobacteria transformation, floral dip, and T_1_ screening were the same as described above. The motif deletion constructs were transformed into a previously characterized *pCLSY3::CLSY3-3xFLAG*/*clsy3* line^40^ to facilitate CLSY3 ChIP experiments. Deletions were identified in the T_2_ generation using the PCR primers listed in **Supplementary Data 11**.

### Methyl-cutting assays

PCR-based methyl-cutting assays were used to semi-quantitatively measure DNA methylation levels at specific loci in different genetic backgrounds. For each assay, genomic DNA was extracted from stage 12 and younger flower buds with the CTAB method^83^ and 500 μg of DNA was digested with the methylation-sensitive restriction enzyme AluI (Thermo Scientific, Cat# FD0014) at 37_°_C for 15 minutes followed by inactivation at 65_°_C for 5 minutes. The digested DNA was amplified by PCR using the primers and conditions listed in **Supplementary Data 11** and visualized using 1.5% agarose gels.

### Mapping-by-sequencing and fine-mapping

The EMS mutant ES1M5S10328 (*rim22-3*) was crossed to its un-mutagenized Col-0 parental line from the HEM population^79^. Flower buds from 48 F_2_ plants were collected and screened using the methyl-cutting assay to identify homozygous mutants. Genomic DNA from 13 mutants were pooled for library preparation using the NEBNext Ultra II DNA Library prep kit (NEB, Cat# E7645L) for Whole Genome Sequencing (WGS). WGS libraries were also constructed for the Col-0 parental line and a few other DNA-methylation-defect mutants identified from this screen (ES1M5S01067/*nrpd1*, EH1S1B549/*nrpd1*, ES1M5S10336/*nrpe1*, ES1M5S01043/*nrpe1*, ES1M5S10229/*nrpe1*, and ES1M5S10051/*ago4*). WGS data for ES1M5S10052/*ago4*^79^ and ES1M5S10328/*rim22*^84^ were published previously. The libraries were sequenced with the single-end 50 bp kit (SE50) on the Illumina HiSeq 4000 platform.

For each line, the fastq files were trimmed to remove low quality reads using trim_galore (v0.6.0)^85^ with default parameters and then mapped to the Arabidopsis TAIR10 genome using BWA (v0.7.17)^86^ with the following functions (aln and samse) and default options. The resulting SAM files were converted to BAM files using samtools (v1.6)^87^ and redundant reads were removed using the rmdup function of samtools (v1.6)^87^. SNP calling was performed using samtools (v1.6)^87^ with the following options: mpileup -Q 30 -C50 -P Illumina -t DP,DV,INFO/DPR,DP4,SP,DV and -Buf. VCF files were generated from the SNP calls using bcftools (v1.9)^88^. The SNPs were first filtered to keep only those induced by EMS (G-to-A and C-to-T) and then annotated using snpEff (v5.0e)^89^ based on the Arabidopsis TAIR10 annotation. SNPs present in the EMS mutants but not in the Col-0 parental line were considered as potential causal SNPs.

For the ES1M5S10328 (*rim22-3*) mapping-by-sequencing library, the read depth and SNP mutation frequency at the final set of SNPs were extracted from the DP and DP4 columns of the VCF file and the mutation frequency was plotted against the SNP positions using R (https://cran.r-project.org/) resulting in the identification of a QTL region on chromosome 3. To fine-map the causative gene, a larger F_2_ population containing 612 plants was grown in the greenhouse and methyl-cutting assays were used to identify homozygous mutants. SNPs within the QTL region were used to develop CAPS or dCAPS markers to genotype the homozygous mutants (primers listed in **Supplementary Data 11**). This approach narrowed the candidate gene to a 1.03-Mb region that included potentially disruptive SNPs in 5 genes (AT3G14910, AT3G16050, AT3G16480, AT3G17010, and AT3G17370).

### Small RNA (smRNA) library preparation, sequencing, and data analysis

#### smRNA library preparation and sequencing

Total RNA extraction and smRNA enrichment was performed following a previously reported protocol^18^. Briefly, 10 μg of total RNA extracted from flower buds (stage 12 and younger) were used for each smRNA library prep using the NEBNext Small RNA Library Prep Kit (NEB, Cat# E7330L). The smRNA libraries were further purified using an 8% acrylamide gel and size selected (130-160 bp) relative to the O’RangeRuler 10 bp DNA Ladder (Thermo Scientific, Cat# SM1313). The size selected smRNA libraries were pooled for sequencing with single-end 50 bp kit (SE50) on Illumina HiSeq 4000 or Nextseq 2000 platforms.

#### smRNA-seq read mapping and siRNA cluster analyses

For each sample, fastq files were trimmed using cutadapt (v1.18)^90^ with the -a AGATCGGAAGAGC option to remove the adapter sequence and the -m 15 option to discard reads shorter than 15 nt. Clean reads were then mapped to the TAIR10 reference genome using ShortStack (v3.8.5)^91^ allowing one mismatch (--mismatches 1). The --mmap f option was also included to assign multiple-mapped reads to a specific locus using the fractional-seeded algorithm. The resulting bam files were then filtered to keep perfectly mapped reads or reads with a single mismatch at the 3’ end using bamtools (v2.5.1)^92^ together with an in-house script (JSON_findPerfectMatches_and_TerminalMisMatches_v3)^18^ to meet the characteristics of Pol-IV transcription^93^. To facilitate downstream analyses, the filtered bam files were used to generate Tag Directories using the makeTagDirectory function of HOMER(v4.10)^94^ with the following options: -format sam -mis 1 -keepAll. To quantify the expression of smRNAs with specific sizes, the tag directories were split by size (20nt-25nt) using a custom perl script (splitTagDirectoryByLength.dev2.pl)^18^ and all subsequent analyses utilized the Tag Directories corresponding to 24nt siRNAs. Read mapping statistics are summarized in **Supplementary Data 1**. All siRNA analyses were carried out using a previously defined set of 12,939 siRNA clusters that were compiled across multiple tissues^40^. Of these clusters, 133 were previously defined as siren loci^40^ and 715 overlap with previously defined HyperTE loci (n=797)^39^. Since some of the 12,939 clusters overlapped with >1 HyperTE locus, this captured nearly all (n=783) of the 797 loci. The 715 clusters that corresponded to HyperTE loci are listed in **Supplementary Data 4.** The full set of 12,939 clusters, the 133 clusters defined as siren loci, as well as the subsets of clusters dependent on specific *clsy* mutants are included in Zhou *et al.*^40^.

#### Differential expression (DE) analysis at previously defined siRNA clusters

The expression levels of 24nt siRNAs for different samples across the full set of 12,939 clusters^40^ were determined using the annotatePeaks.pl function of HOMER (v4.10)^94^ with the following options: -size given -len 1 and -noadj . DE clusters were identified using the R package DESeq2^95^. For the DESeq2 analyses, all reads mapped to the TAIR10 reference genome were used to estimate library size and smRNA sequencing samples performed in different experiments were treated as a covariate to reduce batch effects (DESeq2^95^). Differentially expressed clusters were identified with fold change (FC) ≥ 2 and false discovery rate (FDR) < 0.01 cutoffs. DE clusters in various mutants were listed in **Supplementary Data 2** and **Supplementary Data 3** and visualized as volcano plots made in R with ggplot2 package^96^.

#### siRNA analysis and visualization

To visualize 24nt siRNAs in IGV, UCSC format track files were made using the makeUCSCfile function of HOMER (v4.10)^94^ with the following options: -fragLength 24 and -norm 10000000. The bedGraph format output files were then converted into tdf format files using igvtools (v2.3.68)^97^ with the default options. For all boxplots except Fig. S4, rpkm-normalized siRNAs levels over the indicated clusters were quantified using the annotatePeaks.pl function of HOMER (v4.10)^94^ with the following options (-size given -len 1 and -fpkm) and were plotted in R using the ggplot2 package^96^. For Fig. S4, rpkm-normalized siRNA levels were obtained as stated above, log_2_-converted, and then plotted in R (https://cran.r-project.org/) using the ggplot2 package^96^. For the heatmaps in Fig. 4, log_2_-transformed foldchange values (log_2_FC) obtained from the DESeq2 analysis were plotted in R using the heatmap.2 function of the gplots package (v3.2.0) and the clusters within the different subgroups were ranked based on their log_2_FC values. Scaled Venn diagrams were made using DeepVenn^98^. For the genomic distribution plots, the Arabidopsis genome was split into 100 kb non-overlapping bins. The numbers of siRNA clusters in each bin were counted with bedtools (v2.25.0)^99^ and plotted in R. The Pie charts in Fig. 4 and Fig. S5 were plotted in R. The Scatter plots in Fig.1 and Fig. S3 were plotted in R using the ggplot2 package^96^. For the siRNA heatmaps and metaplots in Fig. 5, ovule WT smRNA-seq data was published previously^40^ and was downloaded from Gene Expression Omnibus (GEO) under accession number GSE165001. smRNA-seq read mapping was performed as described above and BigWig files were generated from the resulting bam files using the bamCoverage function of deeptools (v3.5.1)^100^ with the following options: --normalizeUsing RPKM. The computeMatrix function of deeptools (v3.5.1)^100^ was used to generate an siRNA expression matrix in 50 bp bins across a 2 kb region centered on the motifs under CLSY3 peaks^40^ with the following options: reference-point -b 1000 -a 1000 and --binSize 50. The siRNA levels from 3 WT samples were averaged, log_2_-transformed, and then clustered by kmeans (kmeans 2; see **Supplementary Data 10**) in R. The plotHeatmap functions of deeptools (v3.5.1)^100^ were used to draw the heatmaps and metaplots with the following option: --sortRegions descend and --sortUsing mean, to allow reordering the motifs based on siRNA levels.

### MethylC-seq library preparation, sequencing, and data analysis

Genomic DNA was extracted from flower buds (stage 12 and younger) using the DNeasy Plant Mini Kit (QIAGEN, Cat# 69014). 2 μg of genomic DNA was fragmented using a Covaris S2 sonicator. The sonicated DNA was size-selected using SpeedBeads (Thermo Scientific #65152105050250) to collect fragments 150-300 bp in length. The DNA ends were repaired using the End-It DNA repair kit (epicenter # ER81050) and purified using SpeedBeads. The end-repaired fragments were adenylated using the polymerase activity of the Klenow Fragment (N-terminal truncation of DNA Polymerase I, NEB # M0212L), purified using SpeedBeads, and indexed using Trueseq Adapters (BIOO Scientific # 511911) and T4 DNA ligase (NEB # M0202). After purification using SpeedBeads, the fragments were subjected to bisulfite conversion using the MethylCode Bisulfite Conversion Kit (Invitrogen, MECOV-50). Bisulfite-converted DNA was amplified by PCR using the KAPA HiFi HotStart DNA polymerase (KAPA BIOSYSTEMS # KK2801). The following program was used for the PCR amplification: 95_°_C for 2 min; 98_°_C for 30 s; 4 cycles of (98_°_C for 15 s, 60_°_C for 30 s, 72_°_C for 4 min); and 72_°_C for 10 minutes. The PCR product was purified using SpeedBeads and pooled for sequencing using the pair-end 50 bp kit (PE50) on the Illumina Nextseq 2000 platform.

Reads were mapped to the TAIR10 reference genome using the bs_seeker2-align.py function of BS-Seeker2 (v2.1.8)^101^ allowing 2 mismatches (-m 2). Redundant reads were removed using the remove_clonal.py function of picard tools (http://broadinstitute. github.io/picard/). The bs_seeker2-call_methylation.py function of BS-Seeker2^101^ was used to call methylation at single cytosine resolution with read coverage greater than 4 (-r 4). Read mapping statistics are summarized in **Supplementary Data 5**. To facilitate DNA methylation analyses and visualization, Tag Directory and BigWig data formats were generated as follows: (1) The CGmap output files from the BS-Seeker2 pipeline were converted to wig files using a previously published perl script (BSseeker2_2_wiggleV2_CG_CHG_CHH_allmC.pl)^40^, (2) the wig files were converted into BigWig files using wigToBigWig (v2.9)^102^ following default options, and (3) Tag Directories were made from the wig files using a previously published perl script (parseWig_noChr.v2.pl)^18^ and the makeTagDirectory function of HOMER (v4.10)^94^.

To identify differentially methylated regions (DMRs), the genome was split into 100 bp non-overlapping bins and the average levels of DNA methylation in the CG, CHG, and CHH contexts were compared between each mutant and three different wild-type controls. Thresholds used to declare significant DMRs are as follows: (1) The 100-bp bin should contain at least 4 cytosines in the specified context with read coverage greater than 4, (2) the methylation difference between mutant and wild-type control must be greater than 0.4, 0.2, or 0.1 for CG, CHG, and CHH contexts, respectively, and adjusted p-value must be less than 0.01, and (3) the DMR should pass the first two criteria when compared to three replicates of the wildtype control. DMRs identified in different mutants are summarized in **Supplementary Data 6**.

For DNA methylation boxplots and scatter plots, the average DNA methylation levels at the indicated DMRs or siRNA clusters were calculated based on the Tag Directory files using the annotatePeaks.pl function of HOMER (v4.10)^94^) with the following options (-size given -len 1 and -ratio) and the plots were generated in R using the ggplot2 package^96^. For all metaplots, the computeMatrix function of deeptools (v3.5.1)^100^ was used to generate a matrix with the average methylation levels across 100 bp bins covering the region of interest and 2 kb flanking regions (scale-region -- regionBodyLength 2000 -b 2000 -a 2000 --binSize 100 --skipZeros) based on the BigWig files and the DNA methylation levels at 100 bp non-overlapping bins were plotted in R using the ggplot2 package^96^. Scaled Venn diagrams of overlapping DMRs were made using DeepVenn^98^. For Fig. 5A, the CHH methylation heatmap was generated from the CHH BigWig file using the computeMatrix function of deeptools (v3.5.1)^100^ with the following options: reference-point -b 1000 -a 1000 and –binSize 100. The corresponding metaplot was generated using the plotHeatmap functions of deeptools (v3.5.1)^100^ with the following option: --sortRegions keep. The two heatmap/metaplot groups and within group orders were determined by the siRNA analysis as described in the **siRNA analysis and visualization** section.

### Molecular cloning and generation of epitope tagged RIM22 lines

The *RIM22* complementation line (*pRIM22::RIM22-3xFLAG/rim22*) was constructed using a three part gateway system^103^. A 1,022 bp promoter region of *RIM22* (*pRIM22*) or its coding region minus the stop codon were PCR amplified from Col-0 genomic DNA and cloned into the pDONRp4p1r^103^ or pDONRp221^103^ vectors, respectively, using the Gateway BP Clonase II Enzyme kit (Invitrogen, Cat#11789020) per manufacturer’s instructions. To assemble the final construct, these two plasmids, along with a previously generated pDONRp2p3 plasmid containing a 3xFLAG tag^40^ were recombined into the pK7m34gw destination vector^103^ using Gateway LR Clonase II (Invitrogen, Cat#11791020). Using Gibson cloning (NEB, Cat# E2611S), the regions corresponding to the predicted DNA binding residues (LKF and REI) within the RIM22 B3 DNA binding domain were mutated to encode AAA residues (*pRIM22::RIM22_B3mut_-3xFLAG/rim22)*. The resulting constructs were transformed into AGLO agrobacteria competent cells and introduced into the *rim22* EMS allele (ES1M5S10328/*rim22-3*) background using the floral dip method^82^. T_1_ seeds were selected on half strength Linsmaier and Skoog medium supplemented with 50 mg/L kanamycin. T_2_ families with 3:1 resistant:non-resistant ratios were moved forward and experiments were conducted with homozygous T_3_ or T_4_ seeds. All primers used for these cloning reactions are listed in **Supplementary Data 11**.

### mRNA sequencing and data analysis

Total RNA was extracted from flower buds (stage 12 and younger) using the Zymo Quick-RNA Miniprep Kit (R1055). 1 μg of DNase I treated total RNA was used for mRNA isolation with the NEBNext Poly(A) mRNA Magnetic Isolation Module (NEB, Cat# E7490L). Library preparation was performed using the NEBNext Ultra II RNA Library Prep Kit (NEB, Cat# E7770L). Pooled libraries were sequenced using a 50 bp single-end kit (SE50) on the Illumina Novaseq X plus platform. Reads were mapped to the TAIR10 reference genome using STAR (v2.7.10a)^104^ allowing 2 mismatches and only uniquely mapped reads with the following options: --alignIntronMin 20 -- alignIntronMax 6000 --outFilterMismatchNmax 2 --outFilterMultimapNmax 1 and -- alignMatesGapMax 7500. Read mapping statistics are summarized in **Supplementary Data 8**. To facilitate expression quantification, Tag Directories were generated from the resulting bam files using the makeTagDirectory function of HOMER (v4.10)^94^ with the following options: -format sam -mis 2 and -unique. mRNA expression was quantified using the analyzeRepeats.pl function of HOMER (v4.10)^94^. A custom gtf file (**Supplementary Data 12**^40^) containing genes, transposons, and repeats was used for quantification with the option -condenseGenes to report the highest expressed transcript, and -noadj to report raw read count. UCSC format track files were generated using the makeUCSCfile function of HOMER (v4.10)^94^ with the following options: -fragLength given -norm 10000000 -style rnaseq and -strand both. Differential gene expression analysis was performed with the raw read count in R using the DESeq2 package ^95^. The total read number for each sample was used to normalize the library size. Differentially expressed genes were identified using the following cutoffs: log_2_(FC) ≥ 1 and FDR < 0.01 for upregulated genes, and log_2_(FC) ≤ -1 and FDR < 0.01 for downregulated genes. The volcano plots were made in R using the ggplot2 package^96^. Differentially expressed genes are summarized in **Supplementary Data 9**.

### Transcription factor binding enrichment analysis

To look for transcription factors that bind the same genomic regions as CLSY3 (n=102 ChIP-seq peaks)^40^, a publicly available DAP-seq dataset from a collection of 473 genes^42^ was utilized. Briefly, 10 kb regions centered on the full set of CLSY3 ChIP-seq peaks^40^ were split into 500 bp bins. Bedtools (v2.25.0)^99^ was used to examine if a given bin contained a DAP-seq peak for each of the transcription factors. The proportion of bins across the 102 CLSY3 ChIP-seq peaks that are also bound by a specific transcription factor was plotted in R. The enrichment of REM12 at CLSY3 peaks was further examined with a permutation test to determine whether the observed overlap occurred by chance alone. Briefly, 102 simulated regions with the same sizes as the CLSY3 peaks were randomly assigned to the Arabidopsis genome and the same binning and counting procedure was performed. This permutation was performed 1,000 times to determine a significance threshold of p < 0.01.

### ChIP-qPCR

For the ChIP-qPCR experiment, the pCLSY3::CLSY3-3xFLAG/*clsy3* line, the CLSY3 motif CRISPR deletion line 3-396, which is in the pCLSY3::CLSY3-3xFLAG/*clsy3* background, and the WT control were used to quantify CLSY3 binding. 1 g of flower buds (stage12 and younger) were collected and ground in liquid nitrogen. The powder was cross linked in 25 mL of Nuclei Isolation Buffer (1M sucrose, 60mM hepes, 5mM KCl, 5 mM MgCl_2_, 5 mM EDTA, 1 μM pepstatin, 1 mM PMSF, protease cocktail) with 1% formaldehyde (Sigma, Cat# F8775) for 20 minutes at room temperature. The isolated chromatin was resuspended in 2 mL Nuclei Lysis Buffer (50mM Tris-HCl, 10mM EDTA, 1% SDS, 1μM pepstatin, 1mM PMSF, protease cocktail) in AFA sonication tubes (Covaris, Cat# 520132) and sonicated into ∼500 bp fragments using the Covaris S2 Sonicator with the following settings: Peak Incident Power 175 W, Duty Factor 20, cycle/burst 200, and duration 5 minutes. 200 μL of the sonicated chromatin was saved as input and the remaining chromatin (∼1.8 mL) was diluted to 5.4 mL using ChIP Dilution Buffer (16.7mM Tris-HCl, 167mM NaCl, 1.2mM EDTA, 1.1%Triton X-100, 1μM pepstatin, 1mM PMSF, protease cocktail). 50 μL of anti-FLAG M2 Magnetic beads (Sigma, Cat# M8823) were added to the diluted chromatin and incubated overnight at 4°C. The beads were then washed sequentially with 1 mL of Low Salt Wash Buffer (150 mM NaCl, 0.1% SDS, 1% TritonX-100, 2 mM EDTA, 20 mM Tris-HCl), 1 mL of High Salt Wash Buffer (500 mM NaCl, 0.1% SDS, 1% TritonX-100, 2 mM EDTA, 20 mM Tris-HCl), 1 mL of LiCl Wash Buffer (250 mM LiCl, 1% NP-40, 1% sodium deoxycholate, 1 mM EDTA, 10 mM Tris-HCl), and 1 mL of TE buffer (10 mM Tris-HCl, 1 mM EDTA). Immunoprecipitated chromatin was eluted using 400 μL of elution buffer (50 mM Tris-HCl, 50 mM NaCl, 2 mM EDTA, 1% SDS) at 65°C for 15 minutes. Crossing linking for both the input and IP samples were reversed in 160 mM NaCl, 80 mM DTT, 80 mM NaHCO_3_ buffer overnight at 65°C. The DNA was isolated using Phenol/Chloroform/IAA (Sigma, Cat# 6805), precipitated by ethanol, and resuspended in 20 μL of H_2_O.

The input and immunoprecipitated DNA samples were diluted 10 and 3 times, respectively, before performing the qPCR experiment. The qPCR reaction mix was as follows: Luna Universal qPCR Master Mix (NEB, Cat# M3003X) 5 μL, DNA 3 μL, forward primer (5 μM) 0.3 μL, reverse primer (5 μM) 0.3 μL, H_2_0 1.4 μL. Two technical replicates were employed for both 3x-FLAG-tagged lines and the non-transgenic WT plants. The following program was used for the qPCR using the BIORAD Real-Time PCR System (CFX Opus 384): 95_°_C for 5 min; 45 cycles of (95_°_C for 5 s, 60_°_C for 20 s + plate read); melt curve 65_°_C to 95_°_C with 0.5_°_C increment in 2 s + plate read. qPCR primers are listed in **Supplemental Table 7**. Standard curves were obtained by applying the above qPCR reactions and programs to a dilution series (1x to 1/125x) of the WT input sample. The output SQ values (concentration relative to the input sample of the standard curve) were used for downstream analysis. The percentage of IP relative to input was calculated as follows: SQ-IP/SQ-input/30 × 100%, in which 30 represents dilution factor. The ChIP-qPCR bar chart in Fig. 5 was plotted in Excel with the error bars representing standard deviation of two technical replicates.

### ChIP-seq and DAP-seq data analysis

The CLSY3 ChIP-seq^40^ and REM12 DAP-seq^42^ data was published previously and was downloaded from GEO under accession numbers GSE165001 and GSE60143. For the CLSY3 ChIP-seq, reads were mapped to the TAIR10 reference genome using bowtie (v1.2.2)^105^ allowing 2 mismatches and only uniquely mapped reads were kept with the following options: -m 1 -p 1 -v 2 --all --best and --strata -S. For the REM12 DAP-seq, reads were trimmed using trim_galore (v0.6.0)^85^ and then mapped to the TAIR10 reference genome using bowtie2 (2.2.8)^106^ with default options. The resulting SAM files were converted into BAM files using samtools (v1.6)^87^ with default options. The bam files were filtered to keep unduplicated and uniquely mapped reads using samtools (v1.6)^87^ with the following options: rmdup -s and view -q 20. Read mapping statistics are summarized in **Supplementary Data 7**. To facilitate ChIP-seq and DAP-seq signal visualization, bam files were converted to BigWig files using the bamCoverage function of deeptools (v3.5.1)^100^ following default options. The CLSY3 ChIP-seq and REM12 DAP-seq heatmaps and metaplots in Fig. 5 were drawn using the computeMatrix (reference-point, -b 1000 -a 1000 --binSize 50, and sortRegions keep) and plotHeatmap (sortRegions no) functions of deeptools (v3.5.1)^100^. The two heatmap/metaplot groups and within group orders were determined by the siRNA analysis as described in the **siRNA analysis and visualization** section.

#### REM/RIM alignment and AlphaFold

The alignment of the B3 DNA binding domains of RAV1, RIM22/REM22, REM20, and REM21 were conducted using the CLC Main Workbench v2.0.1 and rasmol coloring. The RIM22/REM22 structure was predicted using AlphaFold^49,50^.

---

**Figure S1.**
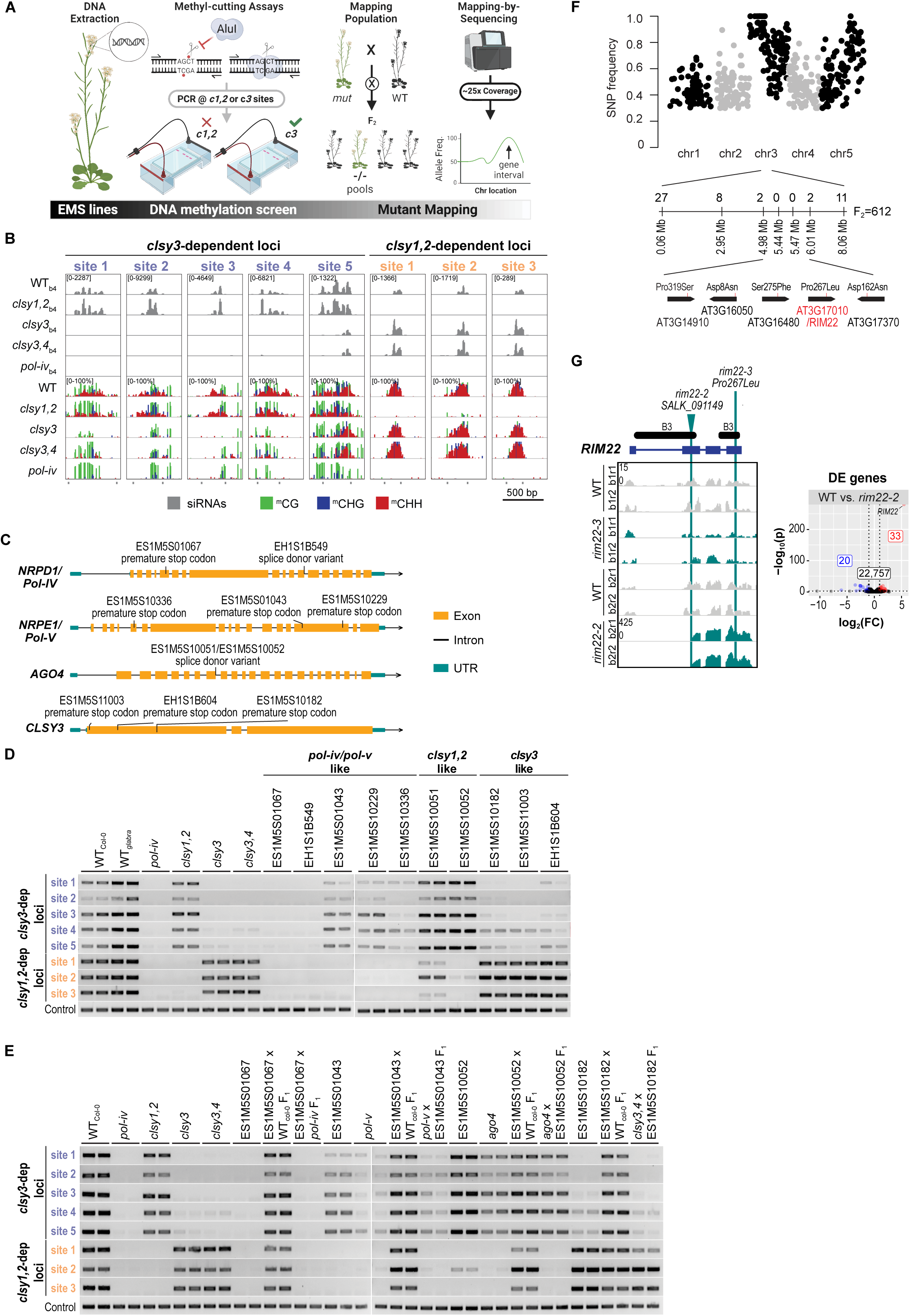
Design of a methyl-cutting screen and identification of DNA methylation mutants. **(A)** Workflow of the methyl-cutting assay-based genetic screen. *Arabidopsis* accessions with EMS-induced mutations were grown in long-day conditions. Flower buds (stage 12 and younger) were collected for genomic DNA extraction, followed by digestion with a methylation-sensitive restriction enzyme, AluI. The digested DNA was amplified by PCR using primers mapping to loci that either lose methylation in *clsy1,2* or *clsy3* mutants. EMS mutants that specifically affect *clsy3*-dependent loci were crossed with the Col-0 parental line, and the F_1_ plants were self-pollinated to generate an F_2_ mapping population. The same methyl-cutting assay was applied to the F_2_ plants and DNA from homozygous mutants were pooled for mapping-by-sequencing. **(B)** Screenshots showing siRNA and DNA methylation levels at *clsy3*- or *clsy1,2*-dependent loci used for the methyl-cutting assay. For each locus, the siRNA expression range is indicated in the upper left corner of the top track and the genotypes, as well as the smRNA-seq batch numbers (b4), are indicated on the left. The DNA methylation tracks are all on a scale of 0-100% and methylation in the CG, CHG, and CHH contexts are shown in green, blue, and red, respectively. The regions amplified for each site are indicated as bars below each set of tracks. DNA methylation data of *pol-iv* was previously published by Zhou *et al*.^18^. **(C)** Gene models showing the mutations identified in known RdDM genes. Vertical black bars indicate SNPs. The mutation type and EMS accession IDs are labeled above each SNP. **(D** and **E)** Gels showing the amplification of DNA at the indicated sites after digestion with the methylation sensitive restriction enzyme, AluI. Two replicates are shown for the genotypes indicated above each set of lanes. The control shows the uniform amplification of a DNA region without the restriction site. All EMS mutants are in the Col-0 background except for EH1S1B549 and EH1SB604, which are in the *glabra* background. In **(D)** EMS mutants were classified into three categories based on phenotype similarity to *pol-iv*/*pol-v*, *clsy1,2*, and *clsy3*. **(F)** Mapping-by-sequencing. EMS-induced SNPs (G to A or C to T) were plotted with the X-axis representing positions and Y-axis representing mutant allele frequency. Fine mapping results are shown as a zoomed-in view of the target region. Vertical bars represent molecular markers, with their chromosomal positions labeled below and the number of recombinants identified between each marker labeled above. The total number of F_2_ plants used for fine mapping is indicated on the right. Genes with mutations in the fine mapping region are displayed, with the SNPs and mutation types labeled above each gene. **(G)** (Left) Genome browser tracks showing the expression levels of *RIM22* in the genotypes indicated on the left. For each genotype, the mRNA-seq batch number (b1 or b2) and replicate number (r1 or r2) are included and the scale (RPK10M) is indicated in the upper left corner of the top track. The locations of the *RIM22* B3 DNA binding domains are shown in black. The positions of the *rim22-2* T-DNA and the *rim22-3* ems allele are represented by a triangle or a vertical bar, respectively. (Right) Volcano plot showing differentially expressed (DE) genes in the *rim22-2* mutant. Samples used for the DEseq analysis are listed in **Supplementary Data 8.** Genes that are downregulated compared to wild-type controls (log_2_FC ≤ -1 and FDR < 0.01) are shown as blue circles, those unaffected are shown as black circles, and those upregulated (log_2_ FC ≥ 1 and FDR < 0.01) are shown as red circles. The number of DE genes in each category are indicated in the correspondingly colored boxes. The red dot corresponding to *RIM22* is marked.

**Figure S2.**
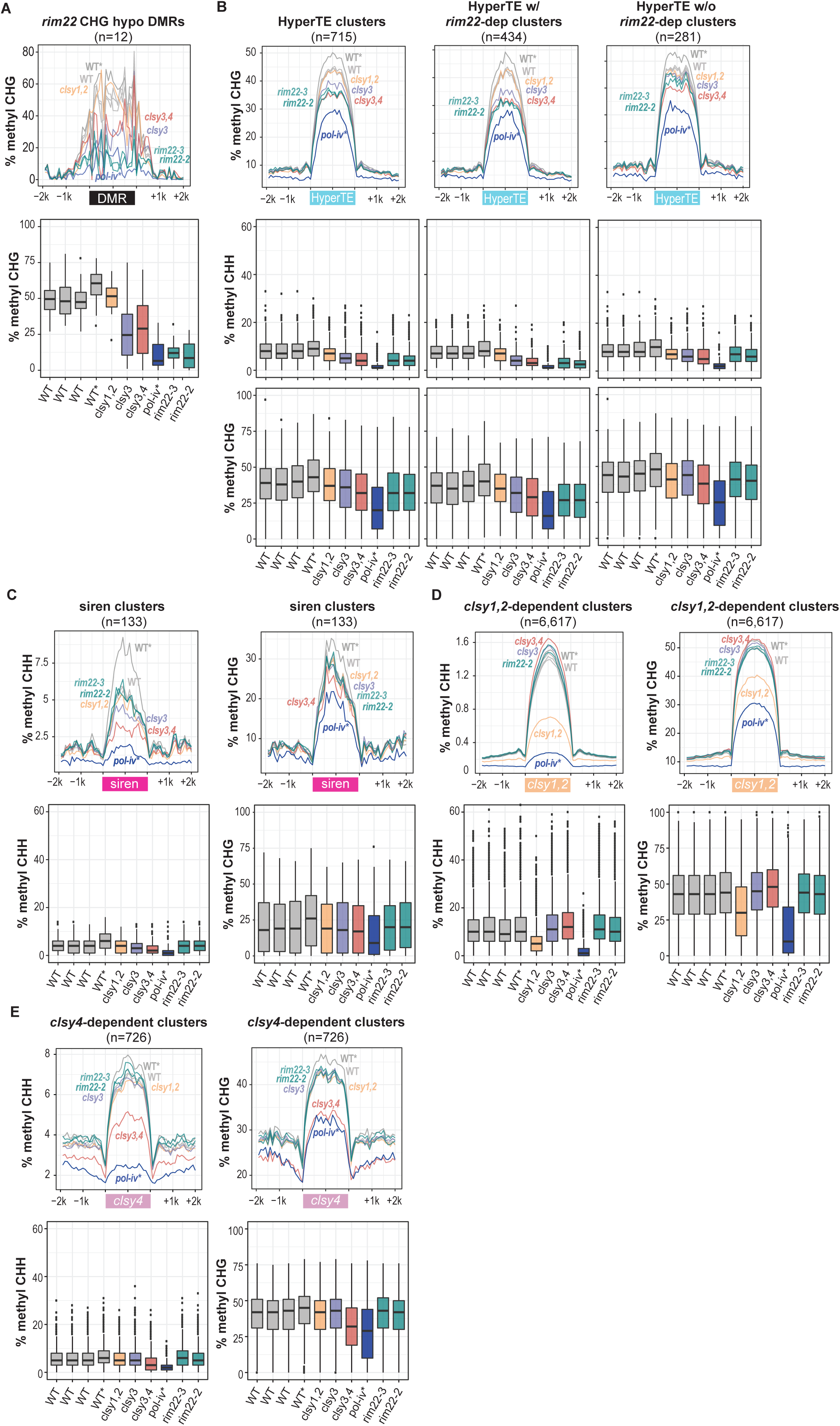
Assessment of DNA methylation levels over *rim22* DMRs, siren loci, and HyperTE loci. **(A-D)** Metaplots (upper) and boxplots (lower) showing the levels of methylation in the indicated contexts at the shared set of CHG DMRs identified in the *rim22* mutants **(A)**, the indicated sets of HyperTE loci **(B)**, siren loci **(C)**, *clsy1,2*-dependent loci **(D)**, or *clsy4*-dependent loci **(E)**. The metaplots show the percent methylation in the indicated genotypes in 100 bp bins across the indicated DMRs and the flanking 2 kb regions. The boxplots show the percent methylation at the indicated DMRs in the genotypes indicated below. For both plots, the two samples from a previous experiment (Zhou *et al.*^18^) are marked with asterisks (*).

**Figure S3.**
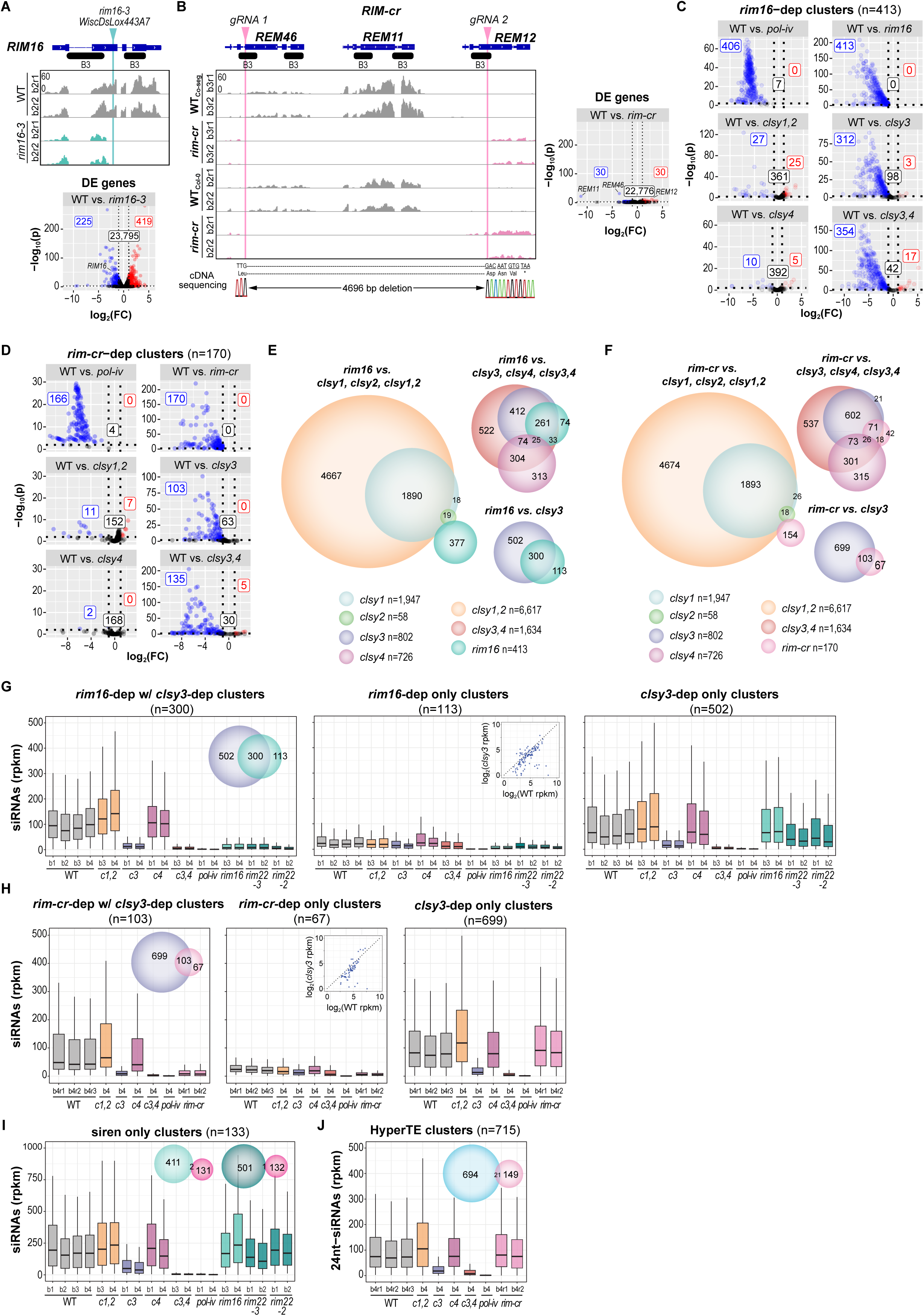
Assessment of the *rim16* and *rim-cr* mutants and their specificity for the regulation of *clsy3*-dependent loci. **(A** and **B)** (Upper) Screenshots showing the expression levels of *RIM16* in **(A)** and *REM46*, *REM11*, and *REM12* in **(B)** with the genotypes, mRNA-seq batches (b2 or b3) and replicates (r1 or r2) indicated on the left. The expression range (RPK10M) for all tracks is indicated in the upper left corner of the top track. For each gene, the locations of the B3 DNA binding domains are shown in black. In **(A)**, the position of the *rim16-3* T-DNA is shown in light teal and in **(B)**, the locations of the two gRNAs used to CRISPR delete the three REM genes are shown in light pink. (Lower or right, respectively) Volcano plots showing differentially expressed (DE) genes in the indicated *rim* mutants. For each plot, clusters that are downregulated compared to wild-type controls (log_2_FC ≤ -1 and FDR < 0.01) are shown as blue circles, those unaffected are shown as black circles, and those upregulated (log_2_ FC ≥ 1 and FDR < 0.01) are shown as red circles. The number of DE genes in each category are indicated in the correspondingly colored boxes. The dots corresponding to *RIM16, REM11, REM12,* and *REM46* are marked. Samples used for the DEseq analysis are listed in **Supplementary Data 8.** In **(B)**, the sequence of the resulting cDNA is included. This sequence shows the generation of a premature stop codon, confirming all three *REM* genes are disrupted. **(C** and **D)** Volcano plots showing siRNA levels at the 413 *rim16*-dependent clusters **(C)** or 170 *rim-cr*-dependent clusters **(D)**. For each plot clusters that are downregulated compared to wild-type (WT) controls (log_2_FC ≤ -1 and FDR < 0.01) are shown as blue circles, those unaffected are shown as black circles, and those upregulated (log_2_ FC ≥ 1 and FDR < 0.01) are shown as red circles. Samples used for the DEseq analysis are listed in **Supplementary Data 1.** The number of clusters in each category are indicated in the correspondingly colored boxes. **(E** and **F)** Scaled Venn diagrams showing the relationships between siRNA clusters reduced in *rim16* or *rim-cr*, respectively, and clusters previously shown to depend on different *clsy* mutants^40^. The circles are colored as indicated below and only overlaps >15 are labeled. **(G** and **H)** Boxplots showing siRNA levels at the indicated clusters. A scaled Venn diagram showing the different categories being compared is shown as an inlay in the first plot. The genotypes, smRNA-seq batches (b1, b2, b3, or b4), and biological replicates (r1, r2, or r3) are indicated below, with each batch representing a distinct biological replicate. Scatter plots are included as inlays in the middle plots showing the expression levels of siRNAs at each of the 113 or 67 loci, respectively, in the *clsy3* mutant compared to its wild-type control. Boxplot outliers were omitted to optimize visualization due to the wide range of data scales. **(I** and **J)** Boxplots showing siRNA levels at the indicated clusters. Scaled Venn diagrams showing the different categories being compared are shown as inlays. The genotypes, smRNA-seq batches (b1, b2, b3, or b4), and biological replicates (r1, r2, or r3) are indicated below, with each batch representing a distinct biological replicate. Boxplot outliers were omitted to optimize visualization due to the wide range of data scales.

**Figure S4.**
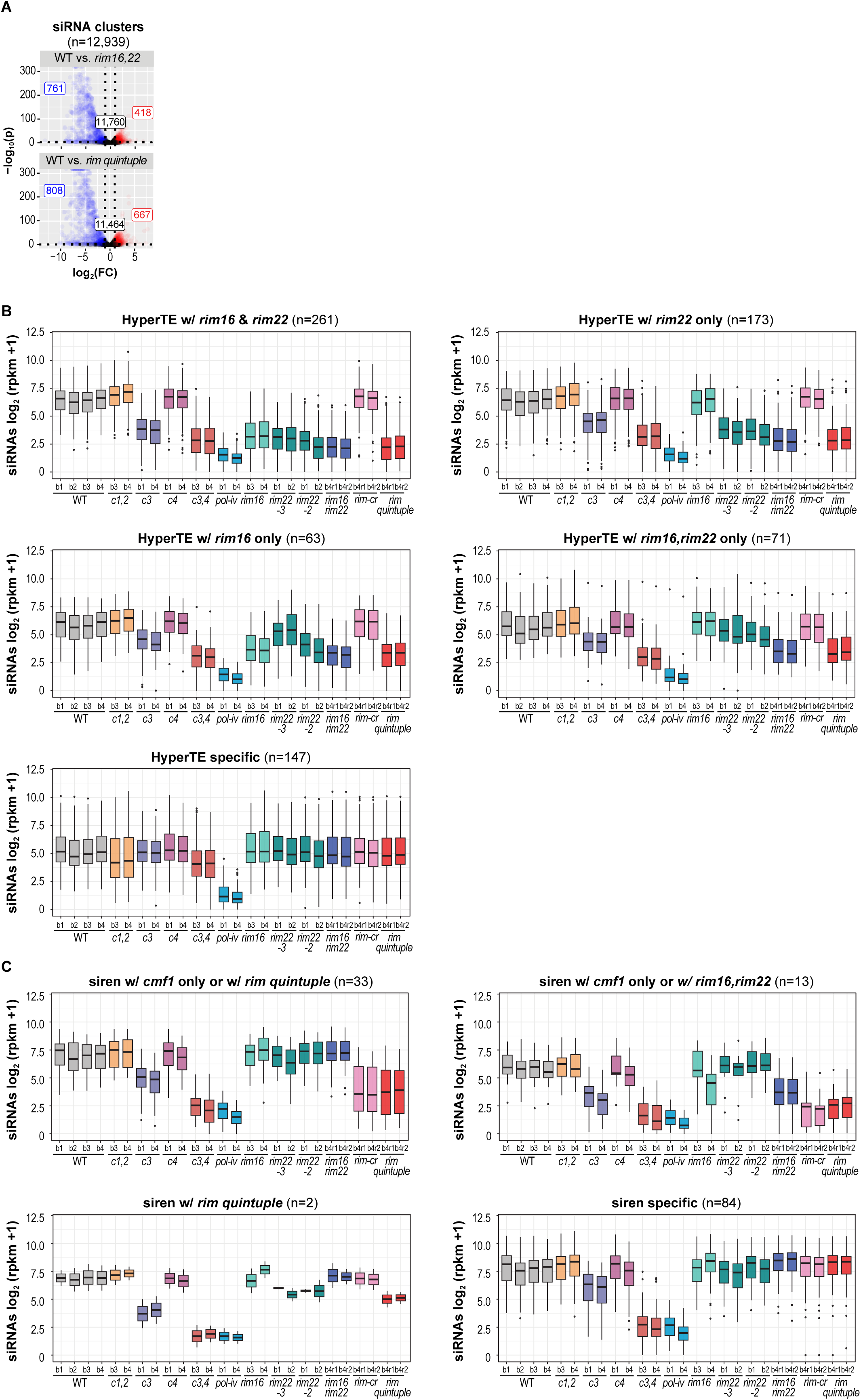
Assessment of siRNA levels in *rim* higher order mutants. **(A)** Volcano plots showing siRNA levels at the full set of RdDM regulated clusters (n=12,939) defined in Zhou *et al.*^40^ in the indicated mutants. For each plot, clusters that are downregulated compared to wild-type (WT) controls (log_2_FC ≤ -1 and FDR < 0.01) are shown as blue circles, those unaffected are shown as black circles, and those upregulated (log_2_ FC ≥ 1 and FDR < 0.01) are shown as red circles. Samples used for the DEseq analysis are listed in **Supplementary Data 1.** The number of clusters in each category are indicated in the correspondingly colored boxes. **(B** and **C)** Boxplots showing siRNA levels at the indicated clusters, which correspond to the heatmap groupings from Fig. 4B and 4D, respectively. The genotypes, smRNA-seq batches (b1, b2, b3, or b4), and biological replicates (r1 or r2) are indicated below, with each batch representing a distinct biological replicate.

**Figure S5.**
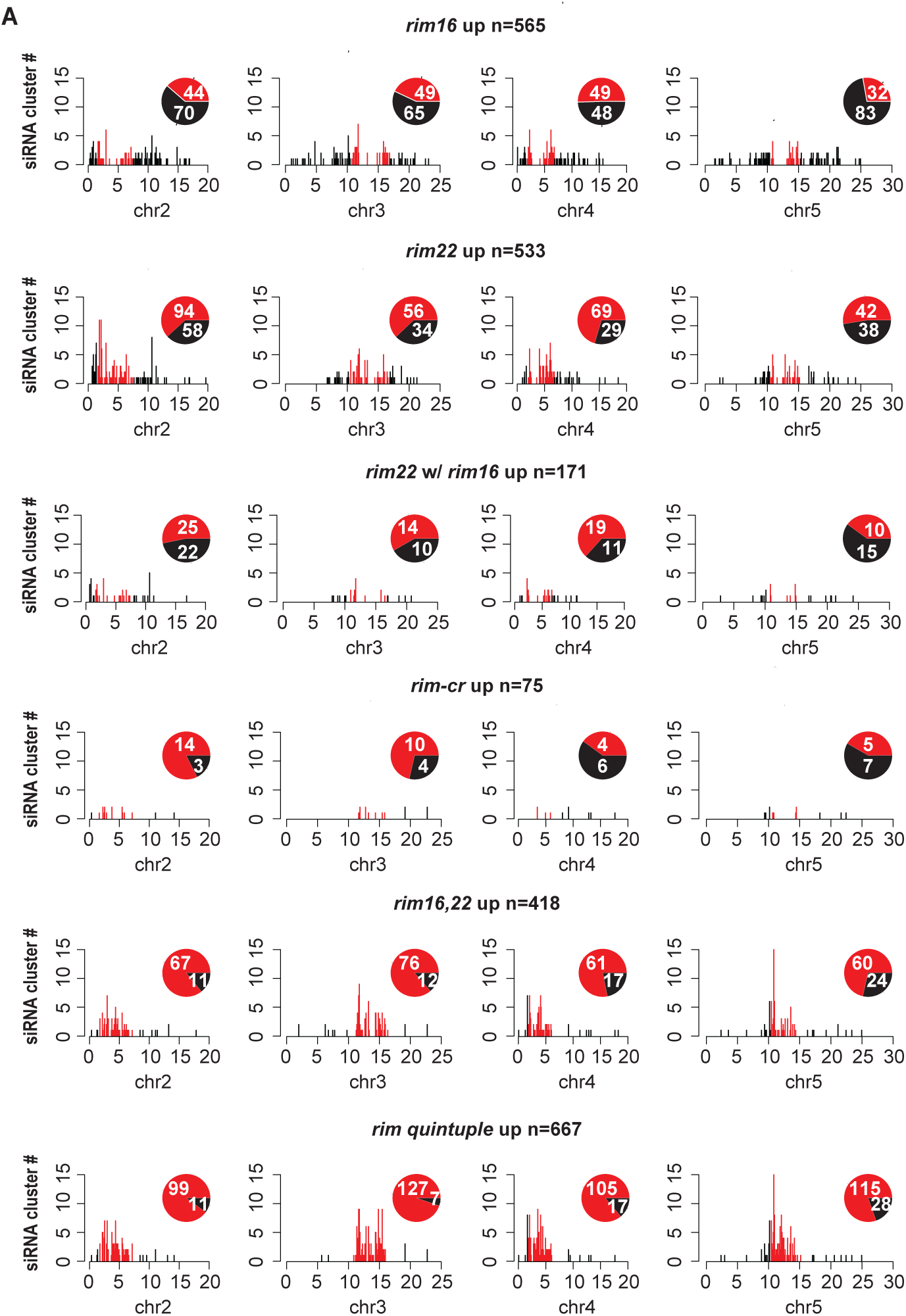
Disruption of multiple RIM TFs results in ectopic siRNA production in pericentromeric heterochromatin. **(A)** Bar and pie charts showing the distributions of upregulated siRNA clusters in the indicated genotypes across chromosomes (chr) 2-5. The distributions across chr1 are shown in Fig. 4E. The total number of clusters across all five chromosomes is indicated above each plot. For both the bar and pie charts, the red and black colors indicate pericentromeric heterochromatin and chromosome arms, respectively. The numbers in the pie charts represent the number of clusters in the corresponding category for each chromosome.

**Figure S6.**
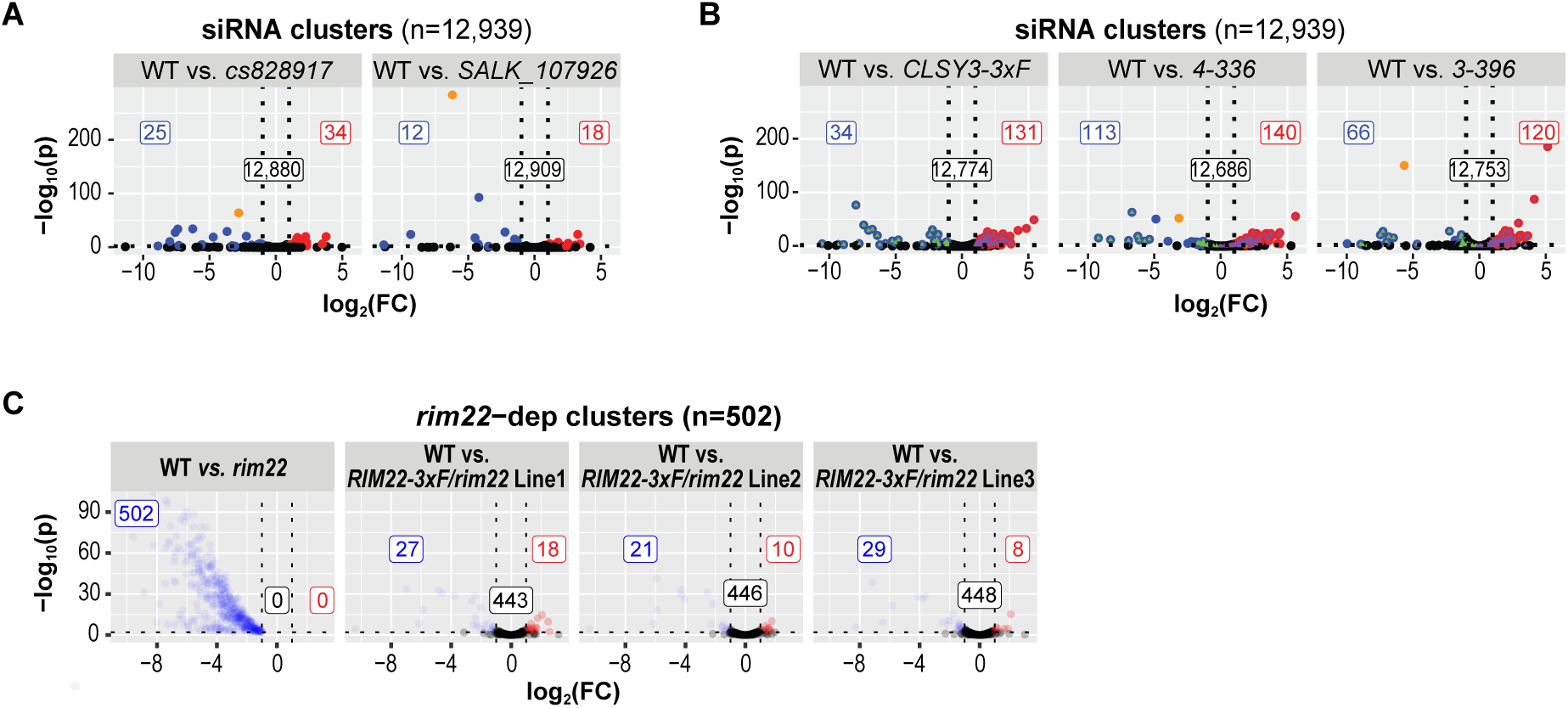
**(A and B)** Volcano plots showing siRNA levels at the full set of RdDM regulated clusters (n=12,939) defined in Zhou *et al.*^40^ in the indicated lines with disrupted CLSY3 motifs. For each plot clusters that are downregulated compared to wild-type (WT) controls (log_2_FC ≤ -1 and FDR < 0.01) are shown as blue circles, those unaffected are shown as black circles, and those upregulated (log_2_FC ≥ 1 and FDR < 0.01) are shown as red circles. Samples used for the DEseq analysis are listed in **Supplementary Data 1.** The number of clusters in each category are indicated in the corresponding colored boxes and the siRNA cluster with the disrupted motif is shown in orange. For **(B)**, the CRISPR editing was conducted in a CLSY3-3xF background (left) and the small number of loci not complemented in this background are marked with triangles in all three volcano plots. **(C)** Volcano plots showing the siRNA defects in *rim22* are complemented by several independent plant lines transformed with a *pRIM22::RIM22-3xFLAG (RIM22-3xF)* construct. For each plot clusters that are downregulated compared to WT controls (log_2_FC ≤ -1 and FDR < 0.01) are shown as blue circles, those unaffected are shown as black circles, and those upregulated (log_2_ FC ≥ 1 and FDR < 0.01) are shown as red circles. Samples used for the DEseq analysis are listed in **Supplementary Data 1.** The number of clusters in each category are indicated in the correspondingly colored boxes.

## Notes

### Competing Interest Statement

The authors have declared no competing interest.

## References

1 Law, J. A. & Jacobsen, S. E. Establishing, maintaining and modifying DNA methylation patterns in plants and animals. Nat Rev Genet 11, 204–220, doi:10.1038/nrg2719 (2010).

2 Calarco, J. P. et al. Reprogramming of DNA methylation in pollen guides epigenetic inheritance via small RNA. Cell 151, 194–205, doi:10.1016/j.cell.2012.09.001 (2012).

3 Consortium, E. P. An integrated encyclopedia of DNA elements in the human genome. Nature 489, 57–74, doi:10.1038/nature11247 (2012).

4 Xu, G. & Law, J. A. Loops, crosstalk, and compartmentalization: it takes many layers to regulate DNA methylation. Curr Opin Genet Dev 84, 102147, doi:10.1016/j.gde.2023.102147 (2024).

5 Martins, L. M. & Law, J. A. Moving targets: Mechanisms regulating siRNA production and DNA methylation during plant development. Current Opinion in Plant Biology 75, 102435 (2023).

6 Xu, G. & Law, J. A. Loops, crosstalk, and compartmentalization: it takes many layers to regulate DNA methylation. Current Opinion in Genetics & Development 84, 102147 (2024).

7 Zhang, H., Lang, Z. & Zhu, J.-K. Dynamics and function of DNA methylation in plants. Nature reviews Molecular cell biology 19, 489–506 (2018).

8 Rymen, B., Ferrafiat, L. & Blevins, T. Non-coding RNA polymerases that silence transposable elements and reprogram gene expression in plants. Transcription 11, 172–191, doi:10.1080/21541264.2020.1825906 (2020).

9 Matzke, M. A. & Mosher, R. A. RNA-directed DNA methylation: an epigenetic pathway of increasing complexity. Nature reviews. Genetics 15, 394–408, doi:10.1038/nrg3683 (2014).

10 Zhang, H., Gong, Z. & Zhu, J. K. Active DNA demethylation in plants: 20 years of discovery and beyond. J Integr Plant Biol 64, 2217–2239, doi:10.1111/jipb.13423 (2022).

11 Herr, A. J., Jensen, M. B., Dalmay, T. & Baulcombe, D. C. RNA polymerase IV directs silencing of endogenous DNA. Science 308, 118–120 (2005).

12 Okano, M., Bell, D. W., Haber, D. A. & Li, E. DNA methyltransferases Dnmt3a and Dnmt3b are essential for de novo methylation and mammalian development. Cell 99, 247–257, doi:10.1016/s0092-8674(00)81656-6 (1999).

13 Onodera, Y. et al. Plant nuclear RNA polymerase IV mediates siRNA and DNA methylation-dependent heterochromatin formation. Cell 120, 613–622, doi:10.1016/j.cell.2005.02.007 (2005).

14 Pontier, D. Reinforcement of silencing at transposons and highly repeated sequences requires the concerted action of two distinct RNA polymerases IV in Arabidopsis. Genes Dev. 19, 2030–2040 (2005).

15 Wierzbicki, A., Haag, J. & Pikaard, C. Noncoding transcription by RNA polymerase Pol IVb/Pol V mediates transcriptional silencing of overlapping and adjacent genes. Cell 135, 635–648 (2008).

16 Cao, X. & Jacobsen, S. E. Role of the arabidopsis DRM methyltransferases in de novo DNA methylation and gene silencing. Curr Biol 12, 1138–1144, doi:10.1016/s0960-9822(02)00925-9 (2002).

17 Cokus, S. J. et al. Shotgun bisulphite sequencing of the Arabidopsis genome reveals DNA methylation patterning. Nature 452, 215–219, doi:10.1038/nature06745 (2008).

18 Zhou, M., Palanca, A. M. S. & Law, J. A. Locus-specific control of the de novo DNA methylation pathway in Arabidopsis by the CLASSY family. Nat Genet 50, 865–873, doi:10.1038/s41588-018-0115-y (2018).

19 Yang, D. L. et al. Four putative SWI2/SNF2 chromatin remodelers have dual roles in regulating DNA methylation in Arabidopsis. Cell Discov 4, 55, doi:10.1038/s41421-018-0056-8 (2018).

20 Law, J. A. et al. Polymerase IV occupancy at RNA-directed DNA methylation sites requires SHH1. Nature 498, 385–389, doi:10.1038/nature12178 (2013).

21 Zhang, H. et al. DTF1 is a core component of RNA-directed DNA methylation and may assist in the recruitment of Pol IV. Proc Natl Acad Sci U S A 110, 8290–8295, doi:10.1073/pnas.1300585110 (2013).

22 Du, J., Johnson, L. M., Jacobsen, S. E. & Patel, D. J. DNA methylation pathways and their crosstalk with histone methylation. Nature reviews Molecular cell biology 16, 519–532 (2015).

23 Johnson, L. M. et al. The SRA methyl-cytosine-binding domain links DNA and histone methylation. Curr Biol 17, 379–384, doi:10.1016/j.cub.2007.01.009 (2007).

24 Rajakumara, E. et al. A dual flip-out mechanism for 5mC recognition by the Arabidopsis SUVH5 SRA domain and its impact on DNA methylation and H3K9 dimethylation in vivo. Genes Dev 25, 137–152, doi:10.1101/gad.1980311 (2011).

25 Ebbs, M. L. & Bender, J. Locus-specific control of DNA methylation by the Arabidopsis SUVH5 histone methyltransferase. Plant Cell 18, 1166–1176, doi:10.1105/tpc.106.041400 (2006).

26 Jackson, J. P. et al. Dimethylation of histone H3 lysine 9 is a critical mark for DNA methylation and gene silencing in Arabidopsis thaliana. Chromosoma 112, 308–315, doi:10.1007/s00412-004-0275-7 (2004).

27 Jackson, J. P., Lindroth, A. M., Cao, X. & Jacobsen, S. E. Control of CpNpG DNA methylation by the KRYPTONITE histone H3 methyltransferase. Nature 416, 556–560, doi:10.1038/nature731 (2002).

28 Du, J. et al. Dual binding of chromomethylase domains to H3K9me2-containing nucleosomes directs DNA methylation in plants. Cell 151, 167–180, doi:10.1016/j.cell.2012.07.034 (2012).

29 Stroud, H. et al. Non-CG methylation patterns shape the epigenetic landscape in Arabidopsis. Nat Struct Mol Biol 21, 64–72, doi:10.1038/nsmb.2735 (2014).

30 Zemach, A. et al. The Arabidopsis nucleosome remodeler DDM1 allows DNA methyltransferases to access H1-containing heterochromatin. Cell 153, 193–205, doi:10.1016/j.cell.2013.02.033 (2013).

31 Lindroth, A. M. et al. Requirement of CHROMOMETHYLASE3 for maintenance of CpXpG methylation. Science 292, 2077–2080, doi:10.1126/science.1059745 (2001).

32 Bartee, L., Malagnac, F. & Bender, J. Arabidopsis cmt3 chromomethylase mutations block non-CG methylation and silencing of an endogenous gene. Genes Dev 15, 1753–1758, doi:10.1101/gad.905701 (2001).

33 Johnson, L. M., Law, J. A., Khattar, A., Henderson, I. R. & Jacobsen, S. E. SRA-domain proteins required for DRM2-mediated de novo DNA methylation. PLoS Genet 4, e1000280, doi:10.1371/journal.pgen.1000280 (2008).

34 Rothi, M. H., Tsuzuki, M., Sethuraman, S. & Wierzbicki, A. T. Reinforcement of transcriptional silencing by a positive feedback between DNA methylation and non-coding transcription. Nucleic Acids Research 49, 9799–9808 (2021).

35 Choi, J., Lyons, D. B., Kim, M. Y., Moore, J. D. & Zilberman, D. DNA Methylation and Histone H1 Jointly Repress Transposable Elements and Aberrant Intragenic Transcripts. Mol Cell 77, 310–323 e317, doi:10.1016/j.molcel.2019.10.011 (2020).

36 Bourguet, P. et al. The histone variant H2A.W and linker histone H1 co-regulate heterochromatin accessibility and DNA methylation. Nat Commun 12, 2683, doi:10.1038/s41467-021-22993-5 (2021).

37 Choi, J., Lyons, D. B. & Zilberman, D. Histone H1 prevents non-CG methylation-mediated small RNA biogenesis in Arabidopsis heterochromatin. Elife 10, doi:10.7554/eLife.72676 (2021).

38 Tang, K., Lang, Z., Zhang, H. & Zhu, J.-K. The DNA demethylase ROS1 targets genomic regions with distinct chromatin modifications. Nature plants 2, 1–10 (2016).

39 Long, J. et al. Nurse cell--derived small RNAs define paternal epigenetic inheritance in Arabidopsis. Science 373, doi:10.1126/science.abh0556 (2021).

40 Zhou, M. et al. The CLASSY family controls tissue-specific DNA methylation patterns in Arabidopsis. Nat Commun 13, 244, doi:10.1038/s41467-021-27690-x (2022).

41 Grover, J. W. et al. Abundant expression of maternal siRNAs is a conserved feature of seed development. Proc Natl Acad Sci U S A 117, 15305–15315, doi:10.1073/pnas.2001332117 (2020).

42 O’Malley, R. C. et al. Cistrome and Epicistrome Features Shape the Regulatory DNA Landscape. Cell 165, 1280–1292, doi:10.1016/j.cell.2016.04.038 (2016).

43 Capilla-Perez, L. et al. The HEM Lines: A New Library of Homozygous Arabidopsis thaliana EMS Mutants and its Potential to Detect Meiotic Phenotypes. Front Plant Sci 9, 1339, doi:10.3389/fpls.2018.01339 (2018).

44 Carrere, S. et al. A fully sequenced collection of homozygous EMS mutants for forward and reverse genetic screens in Arabidopsis thaliana. Plant J, doi:10.1111/tpj.16954 (2024).

45 Hartwig, B., James, G. V., Konrad, K., Schneeberger, K. & Turck, F. Fast isogenic mapping-by-sequencing of ethyl methanesulfonate-induced mutant bulks. Plant physiology 160, 591–600 (2012).

46 Alonso, J. M. et al. Genome-wide insertional mutagenesis of Arabidopsis thaliana. Science 301, 653–657, doi:10.1126/science.1086391 (2003).

47 Swaminathan, K., Peterson, K. & Jack, T. The plant B3 superfamily. Trends Plant Sci 13, 647–655, doi:10.1016/j.tplants.2008.09.006 (2008).

48 Klepikova, A. V., Kasianov, A. S., Gerasimov, E. S., Logacheva, M. D. & Penin, A. A. A high resolution map of the Arabidopsis thaliana developmental transcriptome based on RNA-seq profiling. Plant J 88, 1058–1070, doi:10.1111/tpj.13312 (2016).

49 Jumper, J. et al. Highly accurate protein structure prediction with AlphaFold. Nature 596, 583–589, doi:10.1038/s41586-021-03819-2 (2021).

50 Varadi, M. et al. AlphaFold Protein Structure Database in 2024: providing structure coverage for over 214 million protein sequences. Nucleic Acids Res 52, D368–D375, doi:10.1093/nar/gkad1011 (2024).

51 Yamasaki, K. et al. Solution structure of the B3 DNA binding domain of the Arabidopsis cold-responsive transcription factor RAV1. Plant Cell 16, 3448–3459, doi:10.1105/tpc.104.026112 (2004).

52 Romanel, E. A., Schrago, C. G., Counago, R. M., Russo, C. A. & Alves-Ferreira, M. Evolution of the B3 DNA binding superfamily: new insights into REM family gene diversification. PLoS One 4, e5791, doi:10.1371/journal.pone.0005791 (2009).

53 Alves-Ferreira, M. et al. Global expression profiling applied to the analysis of Arabidopsis stamen development. Plant Physiol 145, 747–762, doi:10.1104/pp.107.104422 (2007).

54 Gomez-Mena, C., de Folter, S., Costa, M. M., Angenent, G. C. & Sablowski, R. Transcriptional program controlled by the floral homeotic gene AGAMOUS during early organogenesis. Development 132, 429–438, doi:10.1242/dev.01600 (2005).

55 Mantegazza, O. et al. Analysis of the arabidopsis REM gene family predicts functions during flower development. Ann Bot 114, 1507–1515, doi:10.1093/aob/mcu124 (2014).

56 Romanel, E. et al. Reproductive Meristem22 is a unique marker for the early stages of stamen development. Int J Dev Biol 55, 657–664, doi:10.1387/ijdb.113340er (2011).

57 Levy, Y. Y., Mesnage, S., Mylne, J. S., Gendall, A. R. & Dean, C. Multiple roles of Arabidopsis VRN1 in vernalization and flowering time control. Science 297, 243–246, doi:10.1126/science.1072147 (2002).

58 Manrique, S. et al. Assessing the role of REM13, REM34 and REM46 during the transition to the reproductive phase in Arabidopsis thaliana. Plant Mol Biol 112, 179–193, doi:10.1007/s11103-023-01357-1 (2023).

59 Yu, Y. et al. Arabidopsis REM16 acts as a B3 domain transcription factor to promote flowering time via directly binding to the promoters of SOC1 and FT. Plant J 103, 1386–1398, doi:10.1111/tpj.14807 (2020).

60 Richter, R. et al. Floral regulators FLC and SOC1 directly regulate expression of the B3-type transcription factor TARGET OF FLC AND SVP 1 at the Arabidopsis shoot apex via antagonistic chromatin modifications. PLoS Genet 15, e1008065, doi:10.1371/journal.pgen.1008065 (2019).

61 Mendes, M. A. et al. Live and let die: a REM complex promotes fertilization through synergid cell death in Arabidopsis. Development 143, 2780–2790, doi:10.1242/dev.134916 (2016).

62 Matias-Hernandez, L. et al. VERDANDI is a direct target of the MADS domain ovule identity complex and affects embryo sac differentiation in Arabidopsis. Plant Cell 22, 1702–1715, doi:10.1105/tpc.109.068627 (2010).

63 Gomez, M. D. et al. Gibberellins negatively modulate ovule number in plants. Development 145, doi:10.1242/dev.163865 (2018).

64 Caselli, F. et al. REM34 and REM35 Control Female and Male Gametophyte Development in Arabidopsis thaliana. Front Plant Sci 10, 1351, doi:10.3389/fpls.2019.01351 (2019).

65 Grover, J. W. et al. Maternal components of RNA-directed DNA methylation are required for seed development in Brassica rapa. The Plant Journal 94, 575–582 (2018).

66 Erhard Jr, K. F., et al. RNA polymerase IV functions in paramutation in Zea mays. Science 323, 1201–1205 (2009).

67 Parkinson, S. E., Gross, S. M. & Hollick, J. B. Maize sex determination and abaxial leaf fates are canalized by a factor that maintains repressed epigenetic states. Developmental biology 308, 462–473 (2007).

68 Pal, A. K., Gandhivel, V. H.-S., Nambiar, A. B. & Shivaprasad, P. Upstream regulator of genomic imprinting in rice endosperm is a small RNA-associated chromatin remodeler. Nature Communications 15, 7807 (2024).

69 Moritoh, S. et al. Targeted disruption of an orthologue of DOMAINS REARRANGED METHYLASE 2, OsDRM2, impairs the growth of rice plants by abnormal DNA methylation. The Plant Journal 71, 85–98 (2012).

70 Wei, L. et al. Dicer-like 3 produces transposable element-associated 24-nt siRNAs that control agricultural traits in rice. Proceedings of the National Academy of Sciences 111, 3877–3882 (2014).

71 Gouil, Q. & Baulcombe, D. C. DNA methylation signatures of the plant chromomethyltransferases. PLoS genetics 12, e1006526 (2016).

72 Felgines, L. et al. CLSY docking to Pol IV requires a conserved domain critical for small RNA biogenesis and transposon silencing. bioRxiv, 2023.2012.2026.573199, doi:10.1101/2023.12.26.573199 (2023).

73 Law, J. A., Vashisht, A. A., Wohlschlegel, J. A. & Jacobsen, S. E. SHH1, a homeodomain protein required for DNA methylation, as well as RDR2, RDM4, and chromatin remodeling factors, associate with RNA polymerase IV. PLoS Genet 7, e1002195, doi:10.1371/journal.pgen.1002195 (2011).

74 Pontier, D. et al. Reinforcement of silencing at transposons and highly repeated sequences requires the concerted action of two distinct RNA polymerases IV in Arabidopsis. Genes & development 19, 2030–2040 (2005).

75 Stroud, H., Greenberg, M. V., Feng, S., Bernatavichute, Y. V. & Jacobsen, S. E. Comprehensive analysis of silencing mutants reveals complex regulation of the Arabidopsis methylome. Cell 152, 352–364 (2013).

76 Dunoyer, P. et al. An endogenous, systemic RNAi pathway in plants. EMBO J 29, 1699–1712, doi:10.1038/emboj.2010.65 (2010).

77 Woody, S. T., Austin-Phillips, S., Amasino, R. M. & Krysan, P. J. The WiscDsLox T-DNA collection: an arabidopsis community resource generated by using an improved high-throughput T-DNA sequencing pipeline. J Plant Res 120, 157–165, doi:10.1007/s10265-006-0048-x (2007).

78 Sessions, A. et al. A high-throughput Arabidopsis reverse genetics system. Plant Cell 14, 2985–2994 (2002).

79 Capilla-Perez, L. et al. The HEM lines: a new library of homozygous Arabidopsis thaliana EMS mutants and its potential to detect meiotic phenotypes. Frontiers in Plant Science 9, 1339 (2018).

80 Wang, Z.-P. et al. Egg cell-specific promoter-controlled CRISPR/Cas9 efficiently generates homozygous mutants for multiple target genes in Arabidopsis in a single generation. Genome biology 16, 1–12 (2015).

81 Xing, H.-L. et al. A CRISPR/Cas9 toolkit for multiplex genome editing in plants. BMC plant biology 14, 1–12 (2014).

82 Clough, S. J. & Bent, A. F. Floral dip: a simplified method for Agrobacterium-mediated transformation of Arabidopsis thaliana. Plant J 16, 735–743 (1998).

83 Doyle, J. J. & Doyle, J. L. A rapid DNA isolation procedure for small quantities of fresh leaf tissue. Phytochemical bulletin (1987).

84 Carrère, S. et al. A fully sequenced collection of homozygous EMS mutants for forward and reverse genetic screens in Arabidopsis thaliana. The Plant Journal (2023).

85 Andrews, S. (Cambridge, United Kingdom, 2010).

86 Li, H. & Durbin, R. Fast and accurate short read alignment with Burrows–Wheeler transform. bioinformatics 25, 1754–1760 (2009).

87 Li, H. et al. The sequence alignment/map format and SAMtools. bioinformatics 25, 2078–2079 (2009).

88 Li, H. A statistical framework for SNP calling, mutation discovery, association mapping and population genetical parameter estimation from sequencing data. Bioinformatics 27, 2987–2993 (2011).

89 Cingolani, P. et al. A program for annotating and predicting the effects of single nucleotide polymorphisms, SnpEff: SNPs in the genome of Drosophila melanogaster strain w1118; iso-2; iso-3. fly 6, 80–92 (2012).

90 Martin, M. Cutadapt removes adapter sequences from high-throughput sequencing reads. EMBnet. journal 17, 10–12 (2011).

91 Johnson, N. R., Yeoh, J. M., Coruh, C. & Axtell, M. J. Improved Placement of Multi-mapping Small RNAs. G3 (Bethesda) 6, 2103–2111, doi:10.1534/g3.116.030452 (2016).

92 Barnett, D. W., Garrison, E. K., Quinlan, A. R., Strömberg, M. P. & Marth, G. T. BamTools: a C++ API and toolkit for analyzing and managing BAM files. Bioinformatics 27, 1691–1692 (2011).

93 Zhai, J. et al. A One Precursor One siRNA Model for Pol IV-Dependent siRNA Biogenesis. Cell 163, 445–455, doi:10.1016/j.cell.2015.09.032 (2015).

94 Heinz, S. et al. Simple combinations of lineage-determining transcription factors prime cis-regulatory elements required for macrophage and B cell identities. Molecular cell 38, 576–589 (2010).

95 Love, M. I., Huber, W. & Anders, S. Moderated estimation of fold change and dispersion for RNA-seq data with DESeq2. Genome biology 15, 1–21 (2014).

96 Villanueva, R. A. M. & Chen, Z. J. (Taylor & Francis, 2019).

97 Wang, Y. et al. ZMP recruits and excludes Pol IV–mediated DNA methylation in a site-specific manner. Science Advances 8, eadc9454 (2022).

98 Hulsen, T. DeepVenn--a web application for the creation of area-proportional Venn diagrams using the deep learning framework Tensorflow. js. arXiv preprint arXiv:2210.04597 (2022).

99 Quinlan, A. R. & Hall, I. M. BEDTools: a flexible suite of utilities for comparing genomic features. Bioinformatics 26, 841–842 (2010).

100 Ramírez, F. et al. deepTools2: a next generation web server for deep-sequencing data analysis. Nucleic acids research 44, W160 (2016).

101 Guo, W. et al. BS-Seeker2: a versatile aligning pipeline for bisulfite sequencing data. BMC genomics 14, 1–8 (2013).

102 Kent, W. J. et al. The human genome browser at UCSC. Genome research 12, 996–1006 (2002).

103 Karimi, M., Depicker, A. & Hilson, P. Recombinational cloning with plant gateway vectors. Plant physiology 145, 1144–1154 (2007).

104 Dobin, A. et al. STAR: ultrafast universal RNA-seq aligner. Bioinformatics 29, 15–21 (2013).

105 Langmead, B. Aligning short sequencing reads with Bowtie. Current protocols in bioinformatics 32, 11.17. 11–11.17. 14 (2010).

106 Langmead, B. & Salzberg, S. L. Fast gapped-read alignment with Bowtie 2. Nature methods 9, 357–359 (2012).

107 O’Malley, R. C. et al. Cistrome and epicistrome features shape the regulatory DNA landscape. Cell 165, 1280–1292 (2016).

